# Active Suppression of the Nigrostriatal Pathway during Optogenetic Stimulation Revealed by Simultaneous fPET/fMRI

**DOI:** 10.1101/2023.10.19.556049

**Authors:** Sabrina Haas, Fernando Bravo, Tudor M. Ionescu, Irene Gonzalez-Menendez, Leticia Quintanilla-Martinez, Gina Dunkel, Laura Kuebler, Andreas Hahn, Rupert Lanzenberger, Bettina Weigelin, Gerald Reischl, Bernd J. Pichler, Kristina Herfert

## Abstract

The dopaminergic system is a central component of the brain’s neurobiological framework, governing motor control, reward responses, and playing an essential role in various brain disorders such as Parkinson’s disease and schizophrenia. Within this complex network, the nigrostriatal pathway represents a critical circuit for dopamine transmission from the substantia nigra to the striatum, a connection that is vital to understanding many of the disease-related dysfunctions. However, stand-alone functional magnetic resonance imaging (fMRI) is unable to study the intricate interplay between brain activation and its molecular underpinnings. In our study, the simultaneous use of [^18^F]FDG functional positron emission tomography (fPET)/BOLD-fMRI provided a new insight that allowed us to demonstrate an active suppression of the nigrostriatal activity during optogenetic stimulation via presynaptic autoinhibition. Our *in vivo* observation emphasizes that the observed BOLD signal depression during neuronal stimulation does not correlate with neuronal inactivity, but results from an active suppression of neuronal firing as shown by the high [^18^F]FDG signal increase. This result not only illustrates the potential of simultaneous fPET/fMRI to understand the molecular mechanisms of brain function but also provides a new perspective on how neurotransmitters such as dopamine influence hemodynamic responses in the brain.

## Introduction

The dopaminergic circuitry is instrumental in numerous essential functions within the nervous system, orchestrating processes related to motor control, reward processing, cognitive functions, and emotional regulation. Its dysfunction has been implicated in a variety of neurological and psychiatric disorders, including Parkinson’s disease (PD), schizophrenia, drug abuse and attention deficit hyperactivity syndrome [1]. As one of the principal neuromodulatory systems in the brain, the dopaminergic system is subject of intense studies, and understanding this complex circuitry is pivotal for elucidating the underlying mechanism of these diseases and for developing targeted therapies. Consequently, the accurate characterization of this circuitry, including its biochemical, structural, and functional aspects, has become vital for both, basic research and clinical applications, offering prospects for improved diagnostics and tailored interventions.

*In vivo* imaging technologies have greatly advanced our understanding of neuronal circuits at the whole brain level. Two powerful non-invasive tools that have emerged in this field are positron emission tomography (PET) with [^18^F]fluoro-2-deoxy-D-glucose ([^18^F]FDG) and functional magnetic resonance imaging (fMRI). Blood-oxygen level dependent (BOLD)-fMRI measures brain activation by monitoring changes associated with blood flow, hence indirectly assessing the areas with increased oxygen demands [2, 3]. On the other hand, [^18^F]FDG-PET provides invaluable insights into the metabolic aspects of neuronal activity [4–6] as a significant portion of glucose is involved in signaling processes, including neuronal firing and neurotransmitter recycling [7]. Both techniques have significantly enhanced our understanding of the functioning of the complex dopaminergic system [8, 9], however the intricate dynamics of dopamine might not be fully captured by either technique alone. Recent developments have led to the integration of PET and MRI into hybrid systems offering remarkable opportunities for comprehensive investigations [10–13].

There is a growing body of evidence suggesting a potential decoupling between metabolic and hemodynamic signals using simultaneously acquired PET/fMRI [13–15], implying that conventional imaging methods may fail to provide a comprehensive or completely accurate representation of the neuronal activity within the brain. This raises the fundamental question, whether we can capture the complete picture of dopaminergic activity and its downstream pathways by using stand-alone imaging techniques.

To explore this compelling question, we initiated a study employing optogenetic stimulation, a cutting-edge method for precisely manipulating neuronal activity [16]. We hypothesized that this approach might unveil hidden dimensions of neuronal functioning, including potential active silencing mechanisms, which have been reported by other techniques [17] and that stand-alone imaging methods might overlook. BOLD-fMRI, with its superior spatial and temporal resolution compared to [^18^F]FDG-PET, which has been improved to study task-related brain activation in a single functional PET (fPET) session [18, 19], each provide unique insights. Previous studies combining [^18^F]FDG-PET and BOLD-fMRI in rats during an electrical whisker stimulation paradigm revealed regional overlaps and mismatches in brain activation pattern between the two modalities [20]. However, these data were not simultaneously acquired, as different stimulation paradigms and time points were used for PET and fMRI.

In this paper, we introduce an innovative approach, fusing optogenetic stimulation with fully simultaneous fPET/fMRI measurements in rats. Our findings not only enhance our understanding of inhibitory mechanisms during neuronal activation but also underscore the efficacy of hybrid PET/MR systems in studying brain function. These insights offer a significant contribution to the field, encouraging further exploration and refining our comprehension of the dopaminergic system.

## Material and Methods

### Animals

All animal experiments were conducted in compliance with the European directives on the protection and use of laboratory animals (Council Directive 2010/63/UE), with the German animal protection law and with approval of the official local authorities (*Regierungspräsidium Tübingen*, permit number R 6/17). Male Long-Evans (n = 36) rats were purchased from Charles River Laboratories (Calco, Lecco, Italy). All rats were maintained in our vivarium on a 12:12 hour light-dark cycle and were kept at a room temperature with 40-60% humidity. Rats had free access to a standard diet and tap water.

### Experimental timeline

A simplified time-course of the experimental procedures is shown in **Fig. 1a**. Rats were randomly divided into two groups and an adeno-associated viral (AAV) vector containing either channelrhodopsin-2 (ChR2) (n = 21) or green fluorescent protein (AAV-GFP) rats (n = 15) was injected into the right substantia nigra pars compacta (SNc). 12 weeks post viral vector injection, an optical fiber was implanted above the SNc and simultaneous [^18^F]FDG-fPET/BOLD-fMRI scans were performed. Laser stimulation was started 20 minutes after start of the fPET/fMRI acquisition using a block design with 3 minutes rest between the blocks. Each 10 minutes stimulation block was divided into 60 seconds *on* and 15 seconds *off* stimulation phases. Light frequency within the *on* phases was 20 Hz with a duty cycle of 50% and a resulting pulse duration of 25 ms. After the acquisition an anatomical MRI was acquired, and the rats were subsequently transcardially perfused, and the brains harvested for *in vitro* validation. ChR2-eYFP and eGFP viral vector expression in the striatum and SNc was confirmed by fluorescence microscopy (**Fig. 1b**).

**Fig. 1:**
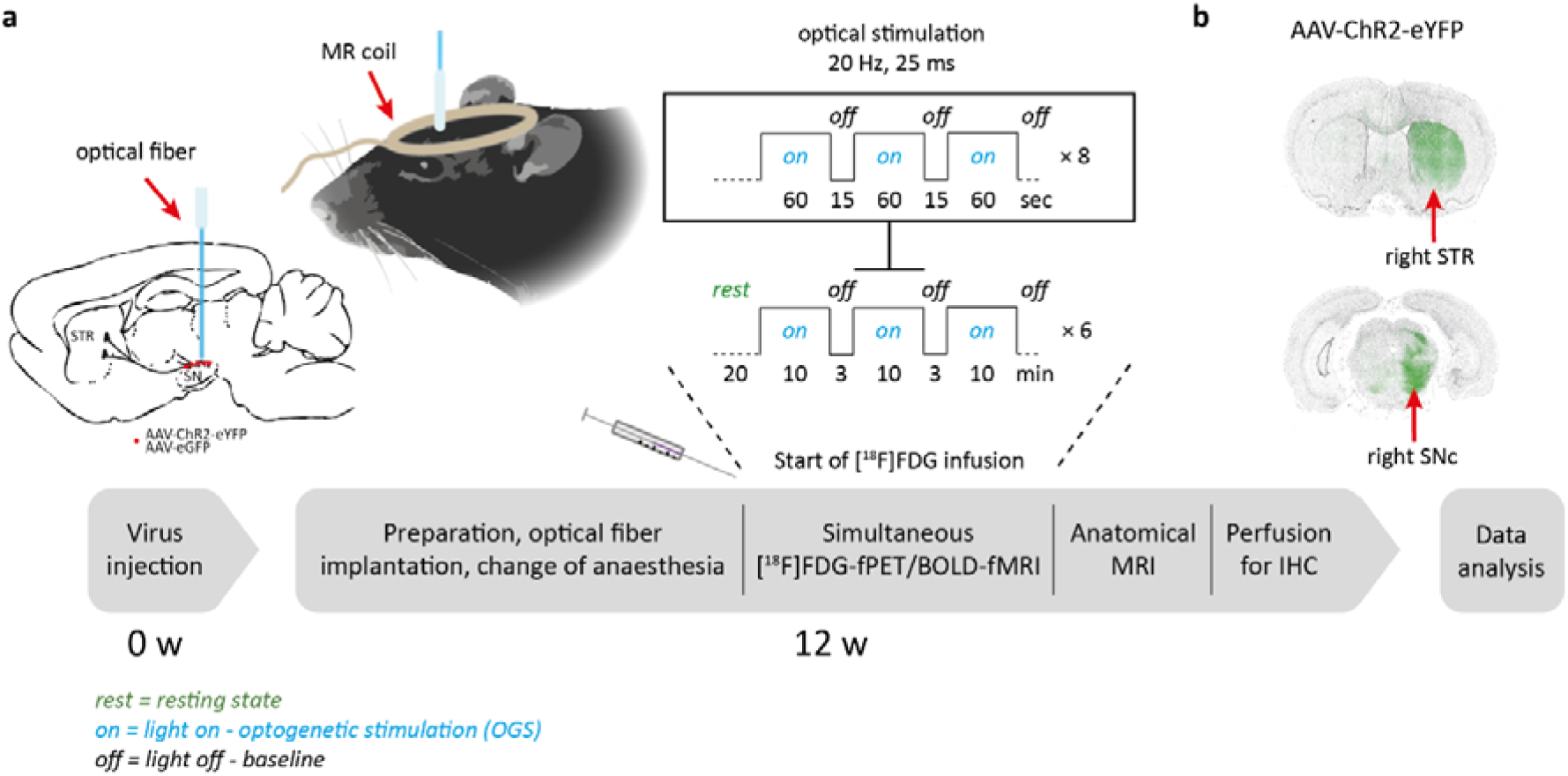
Time course of simultaneous optogenetic [^18^F]FDG-fPET/BOLD-fMRI experiments. (a) AAV-ChR2 or AAV-GFP control virus was injected into the right substantia nigra pars compacta. 12 weeks post viral vector injection, rats were catheterized and intubated. After fiber implantation the [^18^F]FDG-fPET/BOLD-fMRI experiments were acquired using an [^18^F]FDG bolus-infusion protocol during optical stimulation of the substantia nigra pars compacta over 90 minutes using 6 × 10 minute stimulations and 3 minutes rest. Each 10-minute stimulation block consisted of 8 light-on and light-off phases. Within the on phase, a frequency of 20 Hz was set with a duty cycle of 50%, resulting in a pulse duration of 25 ms. After acquisition of an anatomical sequence, the rat brain was transcardially perfused for *in vitro* immunohistochemistry. (b) ChR2 and GFP control virus expression in the striatum and substantia nigra pars compacta was confirmed by fluorescence microscopy of ChR2-eYFP and eGFP. Abbreviations: AAV, adeno-associated virus; BOLD, blood-oxygen level dependent; ChR2, channelrhodopsin-2; eGFP/eYFP, enhanced green/ yellow fluorescent protein; IHC, immunohistochemistry; SNc, substantia nigra pars compacta; STR, striatum

During the time-course of the experiment, 9 rats were excluded from the data analysis due to technical or experimental failures: fiber implantation (n = 3), PET insert (n = 3) and MR (n = 3). During the study, the frequency bandwidth was changed due to a gradient coil exchange: 3 GFP and 5 ChR2 rats were scanned with a frequency bandwidth of 166666.7 Hz; 9 GFP and 13 ChR2 rats were scanned with a frequency bandwidth of 119047.6 Hz.

### Stereotaxic viral vector injection

Rats were allowed to adapt for at least two weeks in the animal facility before viral vector injections. Each rat (n = 36, 375 ± 27 g) was anaesthetized with an intraperitoneal injection of 1 mL/kg of a mixture of fentanyl (0.005 mg/kg), midazolam (2 mg/kg) and medetomidine (0.15 mg/kg). The head was shaved, and the animal placed into a stereotaxic frame. A central incision was made to expose bregma and lambda. A 5 mL Hamilton syringe needle (Hamilton Company, Reno, NV, USA) was enclosed by a glass capillary (inner diameter 50 ± 5 µm, Hilgenberg GmbH, Malsfeld, Germany). Stock solutions of pAAV-hSyn-hChR2(H134R)-EYFP (#26973, AAV5, 1.7×e13 gene copies/mL) or pAAV-hSyn-EGFP (#50465, AAV5, 1.2×e13 gene copies/mL) (Addgene, Inc., Watertown, MA, USA) were diluted to 8.5×e11 gene copies/mL using PBS (Gibco® Dulbecco’s phosphate-buffered saline, Life Technologies, Inc., Carlsbad, CA, USA). 2 µL were slowly injected (0.1 µL every 15 seconds) through a drill-hole into the right SNc (medio-lateral = −2.0 mm, anterior-posterior = −5.0 mm, dorso-ventral = −7.2 mm, according to the stereotaxic atlas of Paxinos and Watson, 1998). To allow for diffusion of the virus into the tissue, the needle was left in place for 5 minutes. Before slowly retracting the needle from the brain (3.5 mm/min), it was withdrawn to −7.0 mm (dorso-ventral) for another 2 minutes. The incision was closed by 4 to 5 stitches and a subcutaneous antidote injection of atipamezol (0.75 mg/kg) and flumazenil (0.2 mg/kg) was administered.

### Simultaneous [^18^F]FDG-fPET/BOLD-fMRI with optogenetic stimulation

#### Optical setup

A 473 nm laser (MBL-III-473nm-100mW, PhotonTec Berlin GmbH, Berlin, Germany) with a maximum output power of 100 mW equipped with a FC/PC fiber coupler having a numerical aperture of 0.22 was used for optogenetic stimulations. The laser was connected (FC/PC MM Fiber Connector, 230 µm, Stainless Steel, Thorlabs, Newton, NJ, USA) to an approximately 6 m long optical fiber (TECS-Clad multimode optical fiber, Thorlabs, Newton, NJ, USA) with a glass fiber core of 200 µm and a numerical aperture of 0.39. The FC/PC connector was assembled and polished in-house using four different polishing sheets sequentially: silicon carbide lapping 5 µm grit, aluminum oxide lapping 3 µm and 1 µm grit, calcinated alumina lapping 0.3 µm grit (Thorlabs, Newton, NJ, USA). The implantable end of the fiber was stripped for at least 2.5 cm and a ceramic ferrule (2.5 mm multimode ceramic ferrule, 231 μm bore size, Thorlabs, Newton, NJ, USA) was glued (LOCTITE® 454™, Henkel AG & Co. KGaA, Dusseldorf, Germany) around the bare fiber end. After drying, the length of the protruding fiber was cleaved to a length of at least 8.2 mm. The laser was coupled to a power supply unit (PSU-III-LED, PhotonTec Berlin GmbH, Berlin, Germany) with TTL modulation up to 1 kHz. It was driven by a stimulus generator (STG 2004, Multi Channel Systems MCS GmbH, Reutlingen, Germany) controlled by a flexible software (MC_Stimulus II, Multi Channel Systems MCS GmbH, Reutlingen, Germany). Fiber output power was measured using a fiber optic power meter (PM20A, Thorlabs, Newton, NJ, USA) prior to each single scan.

#### Animal preparation

AAV-injected rats (n_ChR2_ = 21, n_Ctrl_ = 15, 12 ± 1 weeks post-surgery, 510 ± 36 g) were fasted overnight. Anesthesia was induced with 5% isoflurane evaporated in air in an induction chamber. After loss of the righting reflex, isoflurane was maintained at 2.25-3% evaporated in air at a flow rate of 0.8 L/min. The head was shaved, and a blood sample was collected by puncturing the tail vein for glucose determination (124 ± 13 mg/dL). One tail vein catheter was placed on each side for anesthesia and tracer infusions. Endotracheal intubation was performed using a self-made cannula and an external light source for correct placement of the tube. The small animal ventilator (DC1 73-3629, Harvard Apparatus, Holliston, MA, USA) was set to 60 breaths/min with an inspiration duration of 60% of the ventilation cycle. The end inspiratory pressure was set to approximately 12 cm H_2_O and the flow to 500 mL/min. During preparation and surgery, animals were warmed by a heating pad.

#### Optical fiber implantation

The rat was placed into a stereotaxic frame. A central incision was made to expose bregma and lambda. The optical fiber was inserted through a drilled hole into the right SNc (medio-lateral = −2.0 mm, anterior-posterior = −5.0 mm, dorso-ventral = −7.1 mm, according to the stereotaxic atlas of Paxinos and Watson). Superglue was applied to fixate the ceramic ferrule to the skull. Isoflurane levels were slowly reduced after an initial bolus of 16 mg of α-chloralose (Sigma-Aldrich Chemie GmbH, Taufkirchen, Germany), followed by another bolus containing 5 mg of α-chloralose and 0.25 mg of pancuronium bromide (Inresa Arzneimittel GmbH, Freiburg, Germany). A constant infusion of α-chloralose (20 mg/kg/h) and pancuronium bromide (1 mg/kg/h) was started and maintained during the whole time-course of the experiment along with 0.5% isoflurane evaporated in air.

#### [^18^F]FDG-fPET/BOLD-fMRI

[^18^F]FDG was synthetized using [^18^O]water and the ^18^O(p,n)^18^F nuclear reaction described elsewhere [21]. Simultaneous fPET/fMRI experiments were performed on a small animal 7T MRI system (ClinScan®, Bruker BioSpin MRI GmbH, Ettlingen, Germany) equipped with a small animal PET insert previously described [22]. A linearly polarized RF coil (Bruker BioSpin MRI GmbH, Ettlingen, Germany) with an inner diameter of 72 mm was used for signal excitation and a planar single loop surface coil with an inner diameter of 20 mm (Bruker BioSpin MRI GmbH, Ettlingen, Germany) was used as receiver coil. Rats were placed on a water-heated bed (Medres, Cologne, Germany), connected to the small animal ventilator (DC1 73-3629, Harvard Apparatus, Holliston, MA, USA) and to a feedback temperature control unit (Medres, Cologne, Germany) set to 36.5°C. The temperature was constantly monitored by a rectal probe; oxygen saturation and heartbeat were monitored using a MR compatible pulse oximeter (Bruker BioSpin MRI GmbH, Ettlingen, Germany).

Localizer images were acquired to position the rat brain in the PET/MRI center of the field of view. B0 shimming was performed to optimize magnetic field homogeneity. After an isoflurane wash-out period of at least 1 hour, the PET-insert and a T2*-weighted gradient echo EPI sequence (Duration: 5700 s, TE: 18 ms, TR: 2000 ms, voxel size: 0.27 mm × 0.27 mm × 1.00 mm, FOV: 25 mm × 19 mm, image dimensions: 92 px × 70 px × 20 px, slice thickness: 0.8 mm, slices: 20) covering the brain were started simultaneously. A total of 141 ± 8 MBq [^18^F]FDG were injected 30 seconds after the start of the fPET and fMRI acquisition using a bolus (167 µL/min for 1 minute) plus constant infusion (6.7 µL/min for 93.5 minutes) protocol. Dynamic PET data were acquired for 95 minutes and saved as list-mode files. Laser stimulation was started 20 minutes after start of the simultaneous fPET/fMRI acquisition using a block design described above. Laser irradiance values of 20 ± 3 mW were measured in continuous mode before each fiber implantation using the fiber optic power meter.

At the end of the scan, an anatomical T2 TurboRARE sequence was acquired (TE: 67 ms, TR: 1800 ms, rare factor: 28, averages: 1, FOV: 40 mm × 32 mm × 32 mm, image dimensions: 160 px × 128 px × 128 px, voxel size: 0.25 mm × 0.25 mm × 0.25 mm). To allow for maximal c-fos expression, the animal was transcardially perfused with 50 mL PBS at room temperature, 50 mL PBS cooled to 4°C and 50 mL 4.5% paraformaldehyde (PFA, SAV Liquid Production GmbH, Flintsbach am Inn, Germany) 90 minutes after the start of the first stimulation phase. A second blood sample was collected from an intrathorical vein for glucose determination (86 ± 10 mg/dL) right before perfusion. The brain was surgically removed and fixed in 4.5% formalin (SAV Liquid Production GmbH, Flintsbach am Inn, Germany).

### Imaging data analysis

#### Data preprocessing

fPET list-mode data were divided into 95×1-minute time frames. Sinograms were reconstructed into a dynamic fPET image using OSEM2D reconstruction algorithm. The dynamic brain fPET scans were converted into NIfTI format using PMOD software. fMRI and anatomical images were converted into NIfTI format using Bruker2NIfTI software (v1.0.20170707, Sebastiano Ferraris, University College London).

Data preprocessing was conducted as previously described [13] using Statistical Parametric Mapping 12 (SPM 12, Wellcome Trust Centre for Neuroimaging, University College London, London, United Kingdom) via Matlab (The MathWorks, Natick, MA, USA) and Analysis of Functional NeuroImages (AFNI, National Institute of Mental Health (NIMH), Bethesda, Maryland, USA). In summary, realignment of fMRI and PfET data was performed in SPM. Binary masks were generated from average images and the anatomical MRI scans. With these, the brain was extracted from the fPET, anatomical reference and fMRI image (“skull-stripping”) before co-registration of the fPET and fMRI to the anatomy. Spatial normalization was performed using parameters, which were calculated by comparing the anatomical reference to the Schiffer rat brain atlas [23]. The normalized fMRI and fPET images were smoothed using a 1.5 × 1.5 × 1.5 mm^3^ Gaussian kernel towards the spatial resolution of the PET insert. A temporal high-pass filter with a cut-off frequency of 256 Hz was applied to the fMRI data, with the purpose of removing scanner attributable low frequency drifts in the fMRI time series. Although SPM’s default high-pass cut-off is set to 128 Hz, we increased the cut-off frequency to 256 Hz, since this strategy has been proposed to improve the signal-to-noise ratio when using block lengths of more than 15 seconds off duration as is the case in the present study [24].

Extraction of mean time courses within a region of interest (ROI) was performed using MarsBar [25]. The list of 54 selected ROIs, including abbreviations and volumes is included in the **Supplementary Table 1**.

#### fMRI statistical analysis

Data were analyzed using Statistical Parametric Mapping (SPM), version 12 (www.fil.ion.ucl.ac.uk/spm). A block design was employed for the ChR2 and the GFP groups [24] modeling each of the six 10-minute stimulation blocks using a canonical hemodynamic response function that emulates the early peak at 5 seconds and the subsequent undershoot [26]. The within-subject design matrix for the first level analysis included two regressors: optogenetic stimulation (OGS) and baseline (3 minutes between stimulation blocks). Two contrast images per individual were calculated: OGS > baseline and baseline > OGS.

Between-group approach: Single mean images for each contrast of interest (OGS > baseline and baseline > OGS) were first generated for each subject. Then, a two-sample t-test was carried out to identify the regions that showed significant signal changes between the ChR2 and the GFP groups. Results were thresholded at *p* < 0.001 for voxel-level inference with a cluster-level threshold of *p* < 0.05 corrected for the whole brain volume using family wise error (FWE), which controls for the expected proportion of false-positive clusters.

Within-group approach: Single-subject voxel-wise statistical parametric maps for the aforementioned contrasts were obtained and subjected to group-level one-sample t-tests. The significant map for the group random effects analysis was thresholded at *p* < 0.001 for voxel-level inference with a cluster-level threshold of *p* < 0.05 (FWE corrected).

#### fPET statistical analysis

Between-group approach: [^18^F]FDG-fPET images were first subjected to intensity normalization with reference to the cerebellum [27]. A two-sample t-test was used to compare changes in glucose metabolism induced by optogenetic stimulation during the last 10-minute stimulation block (corresponding to fPET frames 86-95 = time-window with onset at second 2551 with a duration of 600 seconds) between GFP and ChR2 rats. Results were thresholded at *p* < 0.001 for voxel-level inference with a cluster-level threshold of *p* < 0.05 (FWE corrected).

Within-group GLM approach: Modeling of [^18^F]FDG-fPET data with the general linear model (GLM) was done in Matlab as described previously [19, 28]. Here, the GLM is used to separate task effects from baseline by construction of a design matrix that models task effects. This approach is most like conventional fMRI analyses (see within-group fMRI statistics above), thus yielding the term fPET. The design matrix included an OGS regressor and one for the baseline. The OGS regressor was defined as a ramp function with a slope of 1 kBq/frame. The baseline regressor accounts for the continuous uptake of the radioligand due to its irreversible kinetics. It was defined as average of all gray matter voxels, but excluding those voxels declared as activated with the fMRI within-group approach. This approach has been shown to be the best choice in terms of model fits [28], yields comparable results to an independent baseline definition [12]and does not affect test-retest reliability [29]. To increase the SNR of fPET data, a low-pass filter was applied with a cut-off frequency of 5 minutes.

Within-group ICA approach: The data-driven independent component analysis (ICA) approach is a method for recovering underlying signals from linear mixtures of those signals, which draws upon higher-order signal statistics to estimate a set of components that are maximally independent from each other [30]. ICA separates sources by maximizing their non-Gaussianity and, therefore, non-Gaussianity is fundamental for ICA model estimation [31]. One way to understand the connection between independence and non-Gaussianity, is offered by the Central Limit Theorem, which states that the distribution of a sum (or mixture) of random variables tends to be more Gaussian than the original random variables. This, in turn, implies that when the sources are made more non-Gaussian, they become more independent (or unmixed). The distance to a Gaussian can be approximated by using measures of non-Gaussianity, such as skewness and kurtosis, the latter being widely used for estimating non-Gaussianity in ICA. ICA algorithms, including FastICA and Infomax, maximize independence by finding components that have either maximum or minimum kurtosis [32, 33].

ICA has already been applied to [^18^F]FDG-fPET data to investigate brain glucose metabolism and connectivity during task-related designs [34, 35]. We employed the aforementioned strategy [35] and further performed an automatic sorting of the resulting components based upon spatial kurtosis (i.e., spatial sparseness), an approach that proved effective in isolating task-related components without the use of stimulus timing information [36–38]. Task-related components are expected to have non-Gaussian distributions (leading to higher kurtosis values) because they are characterized by transitory, stimulation-induced, increases or decreases in neural activation that are superimposed on a relatively stable background signal [30, 37, 38].

In the case of fPET, ICA first requires a pre-processing step to remove the global baseline signal before the unmixing stage. This technique is conducted to improve the sensitivity for an accurate inference of spatially independent components. Following the procedure described in [34, 35], we first applied whole-brain normalisation to obtain 4D volumes that represented the dynamic relative [^18^F]FDG uptake (time-activity) map [27]. Two further pre-processing steps were implemented before the application of ICA: data reduction and whitening. Data reduction was performed by principal components analysis (PCA) to capture most of the variability in the data (>99%) whilst reducing its dimensionality. Prewhitening was done to improve the convergence of the ICA algorithm and was achieved simultaneously with PCA. To separate the independent components we employed the FastICA algorithm [32, 39, 40]. We estimated twenty components per subject, as this number provided a reasonable trade-off between preserving most of the variance whilst considerably reducing the size of the data. Group-level spatial ICA was conducted using temporal concatenation, which is a widely used approach in group fMRI [39], and which has already been successfully applied to fPET data [34, 35]. The resulting components were sorted according to spatial kurtosis (i.e., a measure of the sparseness of a distribution) following the general framework presented by Lu and Rajapakse 2003 [38]. ICA was implemented with the GIFT v4.0b [39] and CONN v22a [41] toolboxes in MatLab v.R2019a (Natick, Massachusetts USA).

While the GLM approach uses a model-based hypothesis, ICA is data-driven and does not require a priori assumptions on the form and shape of the expected [^18^F]FDG-fPET response. On the other hand, the GLM is simpler to implement and interpret, whereas the ICA approach requires a posteriori selection of components, which can be challenging when the spatial distribution of the effects is unknown. Here, we overcome the need for a manual identification of task effects by automatically sorting components according to spatial kurtosis [37, 38].

#### Percent-overlap-of-activation fPET and fMRI findings

To evaluate the percent-overlap-of-activation between the fPET and fMRI results, we employed the reliability measure proposed by Rombouts *et al.* and Machielsen *et al.*, which is identical to the similarity coefficient proposed by Dice [42–44]. According to this measure, the overlap of activation for any two replications (e.g. *k* and *m*) is established as in **Equation 1**, where *V_k,m_* is the number of voxels identified as activated in both the *k*^th^ and the *m*^th^ replications, and *V_k_* and *V_m_* denote the number of voxels identified as activated in the *m*^th^ and the *k*^th^ experiments, respectively.

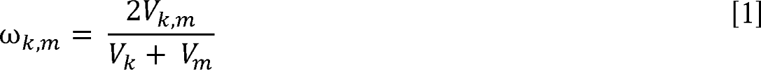

Therefore, ω*_k,m_* is a ratio of the number of voxels identified as activated in both replications to the average number of voxels identified as activated in each replication. It is important to note that this measure spans from 0 (i.e. no overlap) to 1 (perfect overlap) within the identified brain activation.

### Histology

#### Tyrosine-hydroxylase and c-fos immunohistochemistry

Perfused brains were fixated in 4.5% formalin (SAV Liquid Production GmbH, Flintsbach am Inn, Germany) and sectioned into three coronal parts (part A, B and C): one cut was performed approximately through the striatum and the second one through the substantia nigra. Then the tissue was embedded in paraffin. Three rats from each group were selected based on the previous fPET and fMRI results. For histology, 3-5 µm tick sections were cut and stained with hematoxylin and eosin (H&E) and correlated with the “Mouse Brain Atlas” (Allen Reference Atlas – Mouse Brain, available at https://atlas.brain-map.org/) to identify the sections containing the desired anatomical areas (striatum and substantia nigra). Adjacent to those sections, c-fos and tyrosine-hydroxylase (TH) immunohistochemistry (IHC) were performed on an automated immunostainer (Ventana Medical Systems, Inc., Oro Valley, AZ, USA) according to the company’s protocols for open procedures with slight modifications. The slides were stained with the antibodies c-fos (SC-52, Santa Cruz Biotechnology, Dallas, TX, USA) and TH (Tyrosine Hydroxylase (#22941), Immunostar, Hudson, WI, USA). Appropriate positive and negative controls were used to confirm the adequacy of the staining. All samples were scanned with the Ventana DP200 (Roche, Basel, Switzerland) and processed with the Image Viewer MFC Application. Final image preparation was performed with Adobe Photoshop CS6.

The neuronal activation, revealed by c-fos IHC, was bilaterally quantified in the selected rats in the dorsal and ventral striatum, and in the substantia nigra. For this, three ROIs were selected in each target region. See **Supplementary Fig. 1** for more details on the selected ROIs. The number of positive and negative cells was counted at a magnification of 400×. A test for significant differences between right and left ROIs within the group as well as a comparison of the ROIs between GFP and ChR2 expressing rats was performed using a Welch’s t test in Prism 9 (GraphPad Software, LLC, V. 9.3.1, San Diego, CA, USA). No quantification of the TH IHC was performed.

#### GFP and YFP immunofluorescence staining

Adjacent to c-fos, TH and H&E stained sections, a GFP/YFP staining was performed to control for AAV expression. Paraffin sections were rehydrated using a series of xylol and decreasing ethanol concentrations. Antigen retrieval was performed for 15 minutes at 95°C using universal antigen retrieval (R&D Systems, Inc., Minneapolis, MN, USA). Sections were blocked in PBS containing 0.2% Triton-X and 5% bovine serum albumin and stained for GFP or YFP using an anti-GFP antibody (NB100-1614, 1:200, Novus Biologicals, Biotechne, Wiesbaden Nordenstadt, Germany) plus secondary anti-chicken AlexaFluor 555 (A32932, 1:200, Thermo Fisher Scientific Inc., Waltham, MA, USA) together with DAPI (D1306, 1:500, Thermo Fisher Scientific Inc., Waltham, MA, USA). All antibodies were diluted in antibody diluent (IW-1000, IHC World, LLC, Woodstock, MD, USA) and incubated for 1 hour at room temperature. Cover glasses were placed on top using antifade mounting medium (P36980, Thermo Fisher Scientific Inc., Waltham, MA, USA) and sections were acquired on a Leica DMi8 microscope interfaced with Leica LAS X software (Leica Microsystems CMS GmbH, Wetzlar, Germany). The images were further processed with ImageJ.

## Results

### BOLD-fMRI

**Fig. 2a, b** show activated voxels presented as colored t-maps overlaid on an MRI rat brain atlas after between-(n_ChR2_ = 18, n_GFP_ = 12) and within-group (n_ChR2_ = 18) analysis (at threshold *p* < 0.001 voxel-level uncorrected, *p* < 0.05 cluster-level FWE-corrected). Positive hemodynamic responses depicted in red were observed in the right striatum, nucleus accumbens, amygdala, thalamus, substantia nigra and midbrain. Negative responses depicted in green were observed in the left striatum and right and left somatosensory cortex. Higher t-values and spatial extension were obtained using the within-group approach, compared to the between group approach. A list reporting mean t-values and the percentage of activated voxels within a region after cluster-level FWE correction at *p* < 0.05 is shown in **Table 1** for between- and within-group analysis.

**Fig. 2:**
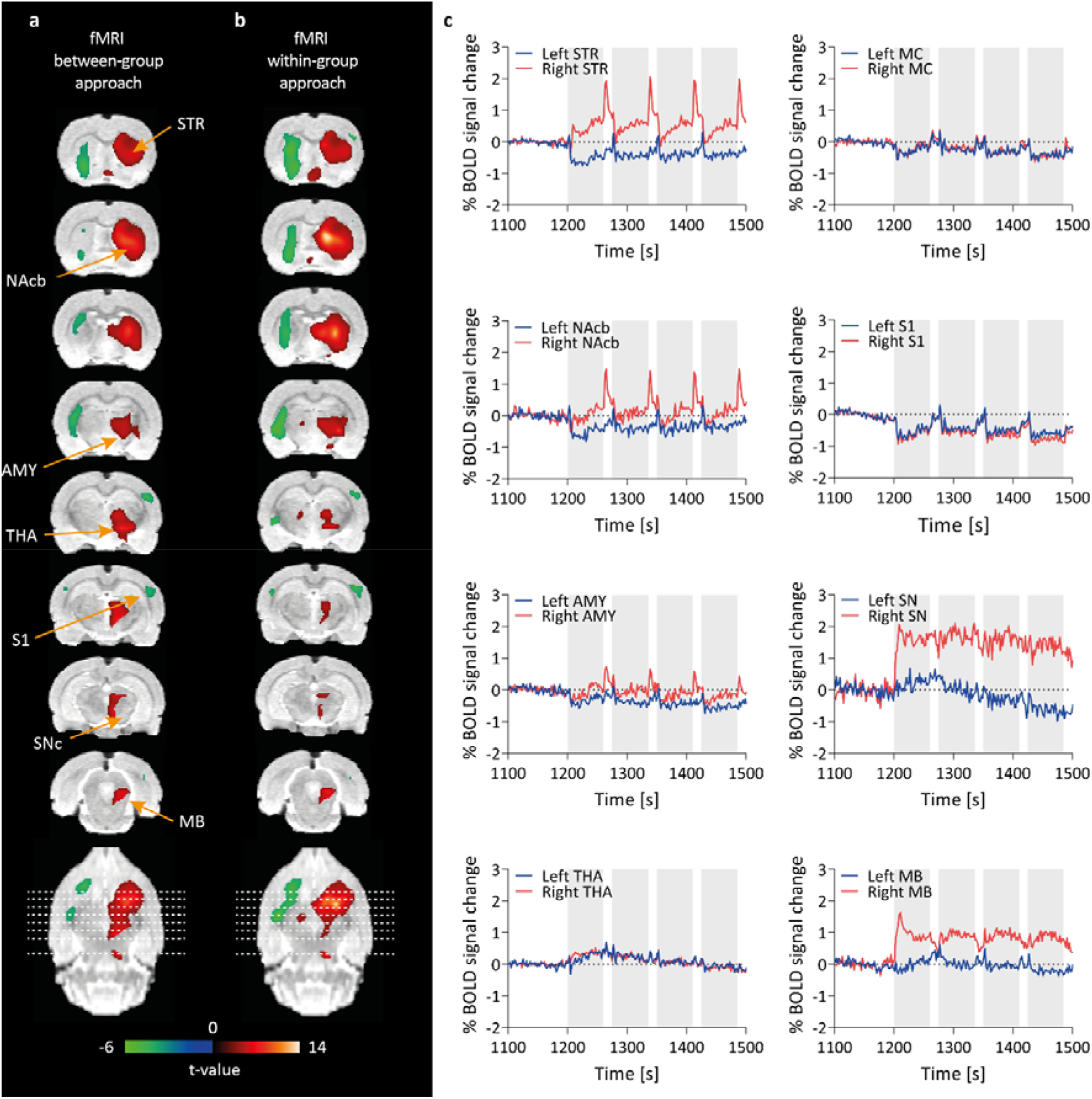
BOLD-fMRI *t*-activation maps after optogenetic SNc stimulation. (a) Between- (ChR2 (n = 18) vs. GFP (n = 12)) and (b) within-group comparison (ChR2, stimulation vs. rest, n = 18) is shown. Positive (red) and negative BOLD responses (green) are shown (FWE-corrected *p* < 0.05 for cluster-level inference). (c) BOLD signal time courses from different brain regions indicate 60 seconds stimulation *on* periods (grey bars). Abbreviations: AMY, amygdala; MB, midbrain; MC, motor cortex; NAcb, nucleus accumbens; S1, somatosensory cortex; SNc, substantia nigra pars compacta; STR, striatum; THA, thalamus

**Table 1:** Percentage of significant voxels per ROI and mean t-values in fMRI.

| Brain region (ROI) | Between-group approach positive BOLD |  | Between-group approach negative BOLD |  | Within-group approach positive BOLD in Chr2 |  | Within-group approach negative BOLD in Chr2 |  |
| --- | --- | --- | --- | --- | --- | --- | --- | --- |
|  | Activated voxels [%] | Mean t | Activated voxels [%] | Mean t | Activated voxels [%] | Mean t | Activated voxels [%] | Mean t |
| R AMY | 2.0 | 3.9 ± 0.3 |  |  | 3.6 | 4.1 ± 0.4 |  |  |
| L AMY |  |  | 0.2 | 3.4 ± 0.02 |  |  | 6.0 | 4.0 ± 0.2 |
| R EC |  |  |  |  | 0.01 | 3.7 ± 0.0 |  |  |
| L EC |  |  | 0.1 | 3.6 ± 0.2 |  |  | 0.4 | 3.9 ± 0.2 |
| L HIP anterior |  |  | 0.9 | 3.6 ± 0.1 |  |  | 0.2 | 3.8 ± 0.1 |
| R HIP posterior | 0.4 | 4.2 ± 0.7 |  |  |  |  |  |  |
| R HYP | 17* | 4.9 ± 1.3 |  |  | 15* | 4.5 ± 0.6 |  |  |
| R INS |  |  |  |  | 0.1 | 3.9 ± 0.1 |  |  |
| L INS |  |  |  |  |  |  | 0.3 | 4.1 ± 0.2 |
| R MB | 3.4* | 4.7 ± 0.6 |  |  | 22* | 5.3 ± 1.0 |  |  |
| L MC |  |  | 0.1 | 3.5 ± 0.04 |  |  | 0.3 | 3.8 ± 0.1 |
| R NAcb | 2.2 | 3.7 ± 0.2 |  |  | 4.0 | 4.1 ± 0.3 |  |  |
| L NAcb |  |  | 18 | 4.0 ± 0.3 |  |  | 18 | 4.4 ± 0.4 |
| L OC |  |  | 0.9 | 3.6 ± 0.2 |  |  |  |  |
| R OFC | 0.2 | 3.6 ± 0.1 |  |  | 0.2 | 4.0 ± 0.2 |  |  |
| L OFC |  |  |  |  |  |  | 1.1 | 4.1 ± 0.2 |
| PAG | 2.4 | 3.7 ± 0.2 |  |  | 2.7 | 4.1 ± 0.4 |  |  |
| R S1 |  |  |  |  | 0.02 | 4.1 ± 0.1 |  |  |
| L S1 |  |  |  |  |  |  | 1.1 | 3.9 ± 0.2 |
| R SC |  |  |  |  | 2.7* | 4.9 ± 0.9 |  |  |
| Sep | 17* | 4.8 ± 1.0 |  |  | 33* | 6.2 ± 2.1 |  |  |
| R SN | 3.6* | 5.0 ± 0.9 |  |  | 2.3 | 4.3 ± 0.4 |  |  |
| R STR | 67* | 5.0 ± 1.2 |  |  | 66* | 5.3 ± 1.5 |  |  |
| L STR |  |  | 31 | 3.9 ± 0.4 |  |  | 58 | 4.3 ± 0.4 |
| R THA | 53* | 4.7 ± 0.7 |  |  | 40* | 5.1 ± 1.2 |  |  |
| L THA |  |  |  |  | 0.9 | 4.3 ± 0.5 |  |  |

Results shown at uncorrected $p < 0.001$ with cluster-level FWE-corrected $p < 0.05$ ; \* markings show areas with significant signal changes at voxel-level FWE-corrected $p < 0.05$ . Abbreviations: AMY, amygdala; Chr2, channelrhodopsin-2; EC, entorhinal cortex; HIP, hippocampus; HYP, hypothalamus; INS, insular cortex; L, left; MB, midbrain; MC, motor cortex; NAcb, nucleus accumbens; OC, olfactory cortex; OFC, orbitofrontal cortex; PAG, periaqueductal gray; R, right; S1, somatosensory cortex; SC, superior colliculus; Sep, septum; SN, substantia nigra; STR, striatum; THA, thalamus

Mean %BOLD signal changes of all ChR2-rats are shown over 400 seconds for selected brain regions (**Fig. 2c**). 60 seconds stimulation blocks are highlighted in grey. Positive BOLD signal changes were observed in the ipsilateral (right) striatum (0.91%), nucleus accumbens (0.64%), amygdala (0.41%), substantia nigra (2.06%) and midbrain (1.59%) during the 60 s stimulation periods (*on* phase). After termination of the stimulation, we observed a BOLD signal overshoot in the ipsilateral (right) striatum (2.06%), nucleus accumbens (1.45%), and amygdala (0.67%), which went back to baseline within the 15 s rest period (*off* phase). Negative BOLD signal changes were observed in the contralateral (left) striatum (−0.74%), nucleus accumbens (−0.84%), amygdala (−0.78%) and in the ipsi- and contralateral motor cortex (−0.63% and −0.66%), and somatosensory cortex S1 (−1.02% and −0.86%). Mean BOLD signal time-courses of all ChR2 rats are shown over the whole scan time for selected regions in **Supplementary Fig. 2**. In GFP control rats, no responses to stimulation were seen in the BOLD signal time-courses (**Supplementary Fig. 3a**).

The BOLD signal time-course of one exemplary ChR2 and GFP rat is plotted over the whole scan time in **Supplementary Fig. 4a**. A BOLD signal increase can be observed in the ipsilateral (right) striatum compared to the contralateral (left) striatum during stimulation (highlighted in grey) in the ChR2 rat, confirming a stimulation-induced BOLD signal change visible on single animal level. No response to stimulation in the ipsi (right)- and contralateral (left) striatum was observed in BOLD signal time courses of the GFP rat.

### [^18^F]FDG-fPET

We applied both, the General Linear Model (GLM) and Independent Component Analysis (ICA), as within-sample methods to examine the optogenetic stimulation experimental data. The optogenetic-stimulation component map appeared as the first ranked component with the highest kurtosis value (9.47), revealing significant [^18^F]FDG uptake in the right substantia nigra, right midbrain, right thalamus, right hypothalamus and right striatum. Additional parameters derived from component’s voxels values distribution, including skewness (a measure of the asymmetry of the distribution), spatial variability (a widespread/clustering measure) and frequency (the centre of mass in spectral power) are shown in **Table 2** (see **Supplementary Table 2** for descriptive measures for all 20 components). Alternatively, the optogenetic-stimulation component can also be identified for its lowest frequency content, following the power spectrum ranking method (Moritz et al., 2002).

**Table 2:** Independent component analysis (ICA). Descriptive measures derived from the independent component’s voxels values distribution.

| Component | Kurtosis | Skewness | Variability | Frequency |
| --- | --- | --- | --- | --- |
| Optogenetic stimulation (via highest kurtosis sorting) | 9.4702 | 1.5296 | 0.89649 | 0.015126 |

Activated voxels are presented as colored t-maps overlaid on an MRI atlas using between-group (**Fig. 3a**), GLM within- (**Fig. 3b**) and ICA within-group (**Fig. 3c**) analysis (n_ChR2_ = 16, n_GFP_ = 14) of [^18^F]FDG-fPET data (at threshold *p* < 0.001 voxel-level uncorrected, *p* < 0.05 cluster-level FWE-corrected). No negative signal changes were observed in the [^18^F]FDG-fPET data. A list reporting mean t-values and the percentage of activated voxels within a region is shown in **Table 3**.

**Fig. 3:**
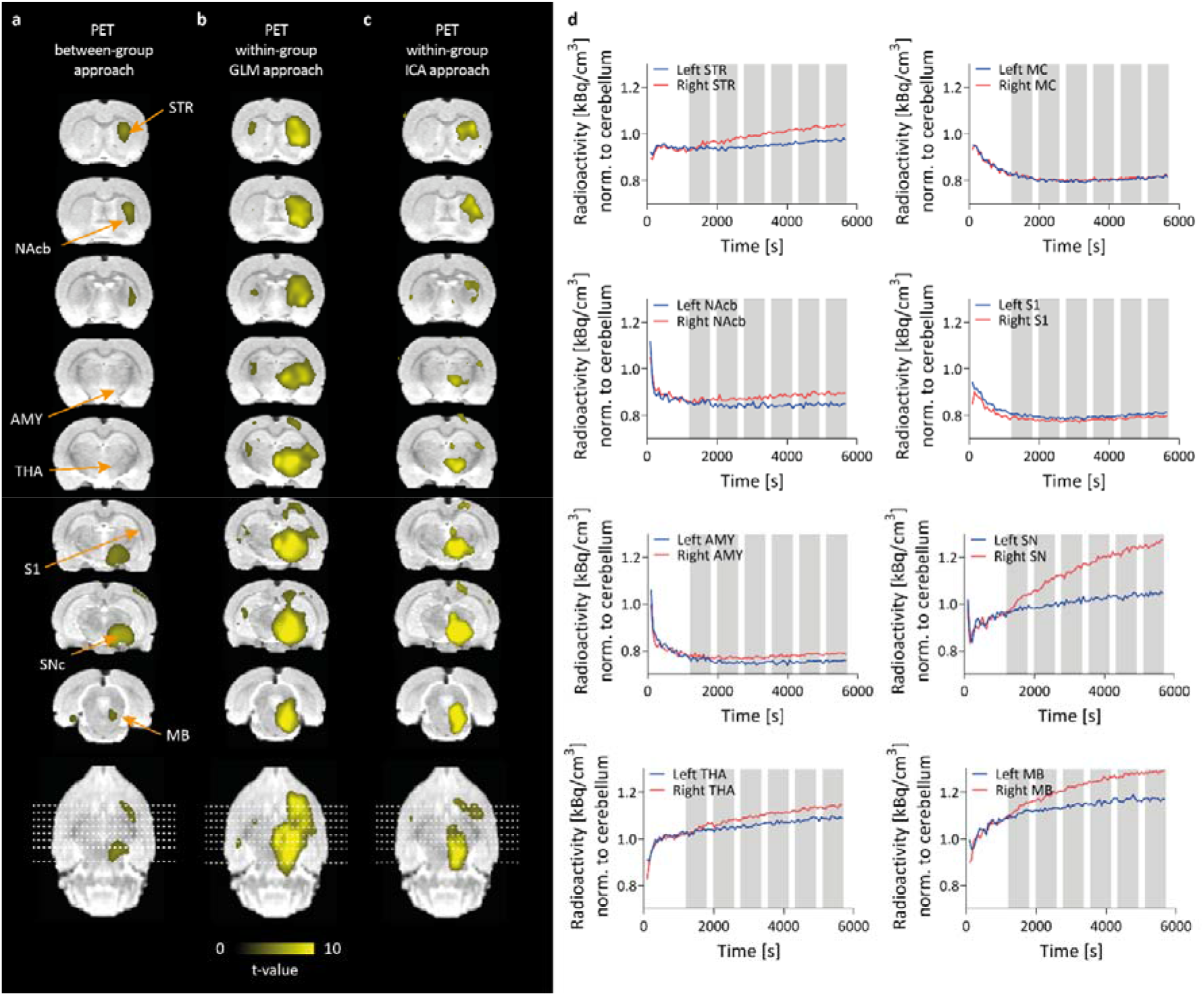
[^18^F]FDG-fPET *t*-activation maps after optogenetic SNc stimulation. (a) Between- (ChR2 (n = 18) vs. GFP (n = 12)), (b) GLM within- (ChR2, stimulation vs. rest, n = 18) and (c) ICA within-group comparison (ChR2, stimulation vs. rest, n = 18) are shown. Positive responses (yellow) are shown (FWE-corrected *p* < 0.05 for cluster-level inference). (d) [^18^F]FDG time activity curves from different brain regions (grey bars indicate 10 minute stimulation blocks). right = ipsilateral, left = contralateral. Abbreviations: AMY, amygdala; GLM, general linear model; ICA, independent component analysis; MB, midbrain; MC, motor cortex; NAcb, nucleus accumbens; S1, somatosensory cortex; SN, substantia nigra; STR, striatum; THA, thalamus

**Table 3:** Percentage of significant voxels per ROI and mean t-values in fPET.

| Brain region (ROI) | Between-group approach |  | Within-group GLM approach |  | Within-group ICA approach |  |
| --- | --- | --- | --- | --- | --- | --- |
|  | Activated voxels [%] | Mean t | Activated voxels [%] | Mean t | Activated voxels [%] | Mean t |
| R AMY | 9.7* | 4.6 ± 0.8 | 11* | 5.0 ± 1.0 | 0.8 | 5.1 ± 0.9 |
| R AUD |  |  | 2.5 | 4.5 ± 0.6 |  |  |
| R EC | 0.5 | 3.9 ± 0.4 | 1.0 | 4.4 ± 0.6 |  |  |
| R HIP anterior |  |  | 5.0 | 4.2 ± 0.4 | 0.7 | 3.9 ± 0.2 |
| L HIP anterior |  |  | 6.1 | 4.1 ± 0.3 |  |  |
| R HIP posterior | 13* | 4.6 ± 0.8 | 34* | 5.6 ± 1.2 | 6.0 | 4.8 ± 0.8 |
| L HIP posterior |  |  | 5.8 | 4.2 ± 0.4 |  |  |
| R HYP | 24* | 4.5 ± 0.7 | 48* | 7.2 ± 2.3 | 36* | 7.5 ± 3.0 |
| L HYP |  |  | 3.4* | 5.4 ± 1.4 | 4.4 | 4.5 ± 0.5 |
| R INS |  |  | 10* | 5.6 ± 1.8 | 2.7 | 4.8 ± 1.0 |
| R MB | 21.2* | 4.4 ± 0.8 | 97* | 7.7 ± 2.1 | 68* | 7.1 ± 2.1 |
| R MC |  |  | 0.6 | 4.1 ± 0.3 |  |  |
| R NAcb |  |  | 12 | 4.9 ± 0.8 | 2.3 | 4.3 ± 0.5 |
| R OFC |  |  | 7.1* | 5.3 ± 1.2 |  |  |
| PG | 3.9 | 3.9 ± 0.3 | 7.7* | 5.2 ± 1.1 | 16 | 4.9 ± 1.0 |
| R PAR |  |  | 29 | 4.3 ± 0.4 |  |  |
| PAG | 0.4 | 3.7 ± 0.2 | 26 | 5.0 ± 0.8 | 9.8* | 5.1 ± 1.0 |
| R RS |  |  | 1.1 | 4.0 ± 0.3 |  |  |
| R S1 |  |  | 3.3 | 4.4 ± 0.5 | 0.5 | 4.4 ± 0.6 |
| R SC |  |  | 7.0 | 4.4 ± 0.6 |  |  |
| Sep |  |  | 0.9 | 4.1 ± 0.3 |  |  |
| R SN | 87* | 5.8 ± 1.0 | 100* | 10.0 ± 1.2 | 98* | 9.8 ± 2.8 |
| L SN |  |  | 11 | 4.7 ± 0.7 | 1.9 | 4.8 ± 0.5 |
| R STR | 27* | 4.3 ± 0.6 | 94* | 6.5 ± 1.5 | 50* | 4.9 ± 0.9 |
| L STR |  |  | 6.6 | 4.2 ± 0.3 |  |  |
| R THA | 0.7 | 3.8 ± 0.3 | 74* | 6.7 ± 2.0 | 24* | 5.3 ± 1.3 |
| L THA |  |  | 4.1*/3.6 | 4.7 ± 0.8/3.9 ± 0.2 | 0.3 | 4.0 ± 0.1 |
| R V1 |  |  | 2.7 | 3.9 ± 0.1 |  |  |

Results shown at uncorrected $p < 0.001$ with cluster-level FWE-corrected $p < 0.05$ ; \* markings show areas with significant signal changes at voxel-level FWE-corrected $p < 0.05$ . Abbreviations: AMY, amygdala; AUD, auditory cortex; ChR2, channelrhodopsin-2; EC, entorhinal cortex; GFP, green fluorescent protein; GLM, general linear model; HIP, hippocampus; HYP, hypothalamus; ICA, independent component analysis; INS, insular cortex; L, left; MB, midbrain; MC, motor cortex; NAcb, nucleus accumbens; OFC, orbitofrontal cortex; PAG, periaqueductal gray; PG, pituitary gland; R, right; RS, retrosplenial cortex; S1, somatosensory cortex; SC, superior colliculus; Sep, septum; SN, substantia nigra; STR, striatum; THA, thalamus; V1, visual cortex

During optogenetic stimulation, the GLM approach yielded increased [^18^F]FDG uptake in similar areas as after using the between- and ICA within-group approach (right substantia nigra, right midbrain, right hypothalamus, and right striatum). Additionally, the GLM strategy showed strong uptake in the right insular cortex, thalamus, right hippocampus posterior, left hypothalamus, right orbitofrontal cortex, pituitary gland, and right amygdala. Overall, the all three approaches successfully identified [^18^F]FDG increases in the right dorsal striatum and the right substantia nigra during the optogenetic stimulation, but the peak and spatial extent of the activations differed between the three methods.

Mean normalized TACs of all ChR2 expressing rats are shown over the whole scan time for selected brain regions. 10 min stimulation blocks are highlighted in grey (**Fig. 3d**). We observed a gradual increase of [^18^F]FDG during the stimulation in the ipsilateral (right) striatum nucleus accumbens, thalamus, substantia nigra and midbrain compared to the contralateral side which started with a delay of 2-5 minutes after start of the stimulation, while little to no changes were observed in the amygdala, motor cortex and somatosensory cortex. Mean normalized TACs of all GFP expressing rats are shown over 95 minutes for selected regions (**Supplementary Fig. 3b**). No changes between the left and right hemisphere of the selected brain regions were observed in the GFP group.

[^18^F]FDG activity of one exemplary ChR2 and GFP rat is plotted over the whole scan time in **Supplementary Fig. 4b**. [^18^F]FDG showed an increased accumulation in the ipsilateral (right) striatum compared to the contralateral (left) striatum in the ChR2 rat, while no differences between the ipsi-(right) and contralateral (left) striatum was found in the GFP rat, confirming a stimulation-induced increase in [^18^F]FDG metabolism.

### [^18^F]FDG-fPET and BOLD-fMRI: Comparison of hemodynamic and metabolic responses to stimulation

**Fig. 4a, b** show a comparison of activated voxels from BOLD-fMRI and [^18^F]FDG-fPET as colored t-maps overlaid on an MRI atlas (between- and within group comparison) (at threshold *p* < 0.001 voxel-level uncorrected, *p* < 0.05 cluster-level FWE-corrected). For within group fPET we show the results from the data-driven independent component analysis approach (ICA with kurtosis/frequency sorting), which does not require a priori information, as is the case for the GLM fPET approach.

**Fig. 4:**
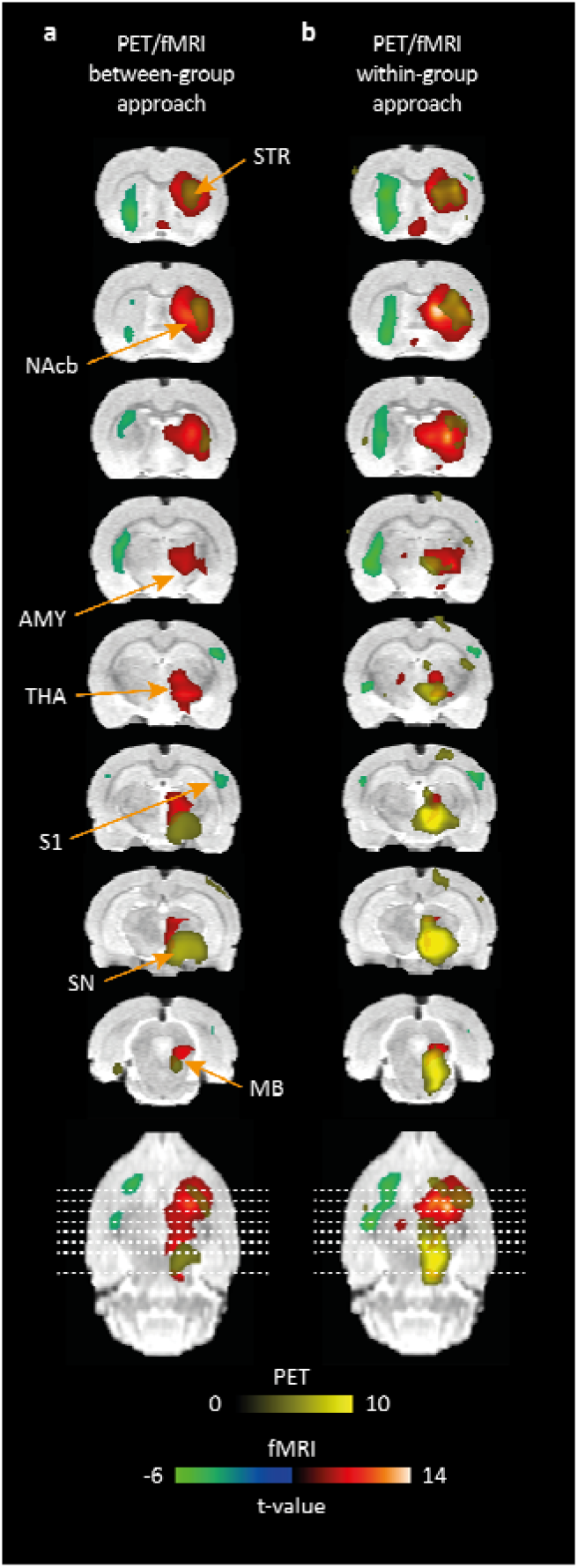
Overlay of [^18^F]FDG-fPET and BOLD-fMRI activation maps after (a) between- and (b) within-group analysis. An overlay of significantly activated and deactivated areas (FWE-corrected *p* < 0.05 for cluster-level inference) after optogenetic stimulation of the substantia nigra pars compacta in fPET and fMRI is shown in colored overlays on a rat brain atlas. Activated areas in fMRI are depicted in red, deactivated areas in fMRI are depicted in green and activated areas in fPET are depicted in yellow. The spatial extension of activated areas between modalities differs. No negative responses were seen in fPET. Abbreviations: AMY, amygdala; ICA, independent component analysis; MB, midbrain; MC, motor cortex; NAcb, nucleus accumbens; S1, somatosensory cortex; SN, substantia nigra; STR, striatum; THA, thalamus

Dice similarity coefficients were calculated in order to quantify overlapping voxels of both modalities, revealing five overlapping regions after between-group fMRI/fPET analysis, namely, the right hypothalamus (D = 0.277), right midbrain (D = 0.165), right substantia nigra (D = 0.026), right striatum (D = 0.054) and right thalamus (D = 0.024) (**Table 4**). Within-group analysis revealed six overlapping regions: right striatum (D = 0.743), right nucleus accumbens (D = 0.444), right insular cortex (D = 0.080), right thalamus (D = 0.221), right somatosensory cortex (D = 0.042) and right hypothalamus (D = 0.018). Largest differences were observed in the spatial extension, most predominantly in the right striatum and right substantia nigra. Independent of the approach, BOLD-fMRI activation maps of the striatum show a larger spatial extension than fPET activation maps, while the opposite was observed in the substantia nigra. This is also confirmed when comparing the % of activated voxels per region presented in **Table 3**. Between-group approach: striatum: 67% vs. 27% and substantia nigra: 3.6% vs. 87%. Within-group approach: striatum: 66% vs. 50% and substantia nigra: 2.3% vs. 98%). Negatively activated areas were only identified in BOLD-fMRI. These were mainly located on the contralateral side, but also on the ipsilateral side to stimulation. Coordinates of peak t-values were extracted from regions of interest activated in both modalities to quantify the distance of the respective activation centers (**Table 4**).

**Table 4:** Distance of t-value peak location between fPET and fMRI and Dice similarity coefficient (between- and within-group analysis)

| Brain region (ROI) | Distance activation centers [mm]<br>BETWEEN | Distance activation centers [mm]<br>WITHIN | Dice similarity coefficient<br>BETWEEN | Dice similarity coefficient<br>WITHIN |
| --- | --- | --- | --- | --- |
| R AMY | 5.0 | 5.0 | - | - |
| R HIP posterior | 2.2 | - | - | - |
| R HYP | 1.6 | 2.1 | 0.277 | 0.018 |
| R MB | 2.1 | 2.7 | 0.165 | 0 |
| PAG | 1.9 | 1.0 | - | - |
| R SN | 1.3 | 0.9 | 0.026 | 0 |
| R S1 | - | 2.8 | - | 0.042 |
| R INS | - | 3.7 | - | 0.080 |
| R NAcb | - | 0.9 | - | 0.444 |
| R STR | 1.2 | 2.9 | 0.548 | 0.743 |
| R THA | 3.5 | - | 0.024 | 0.0221 |

Abbreviations: AMY, amygdala; HIP, hippocampus; HYP, hypothalamus; ICA, independent component analysis; L, left; MB, midbrain; PAG, periaqueductal gray; R, right; SN, substantia nigra; STR, striatum; THA, thalamus; NAcb, nucleus accumbens; INS, insular cortex; S1, somatosensory cortex.

### Validation of brain activation using c-fos immunohistochemistry

Qualitative assessment of c-fos+ staining revealed a higher number of c-fos+ cells in the right dorsal striatum of ChR2 rats compared to the left dorsal striatum and compared to ipsi- and contralateral dorsal striata of GFP rats (see **Fig. 5a**, and **Supplementary Fig. 5**). No differences were observed between the right and left substantia nigra of both groups (**Fig. 5b**).

**Fig. 5:**
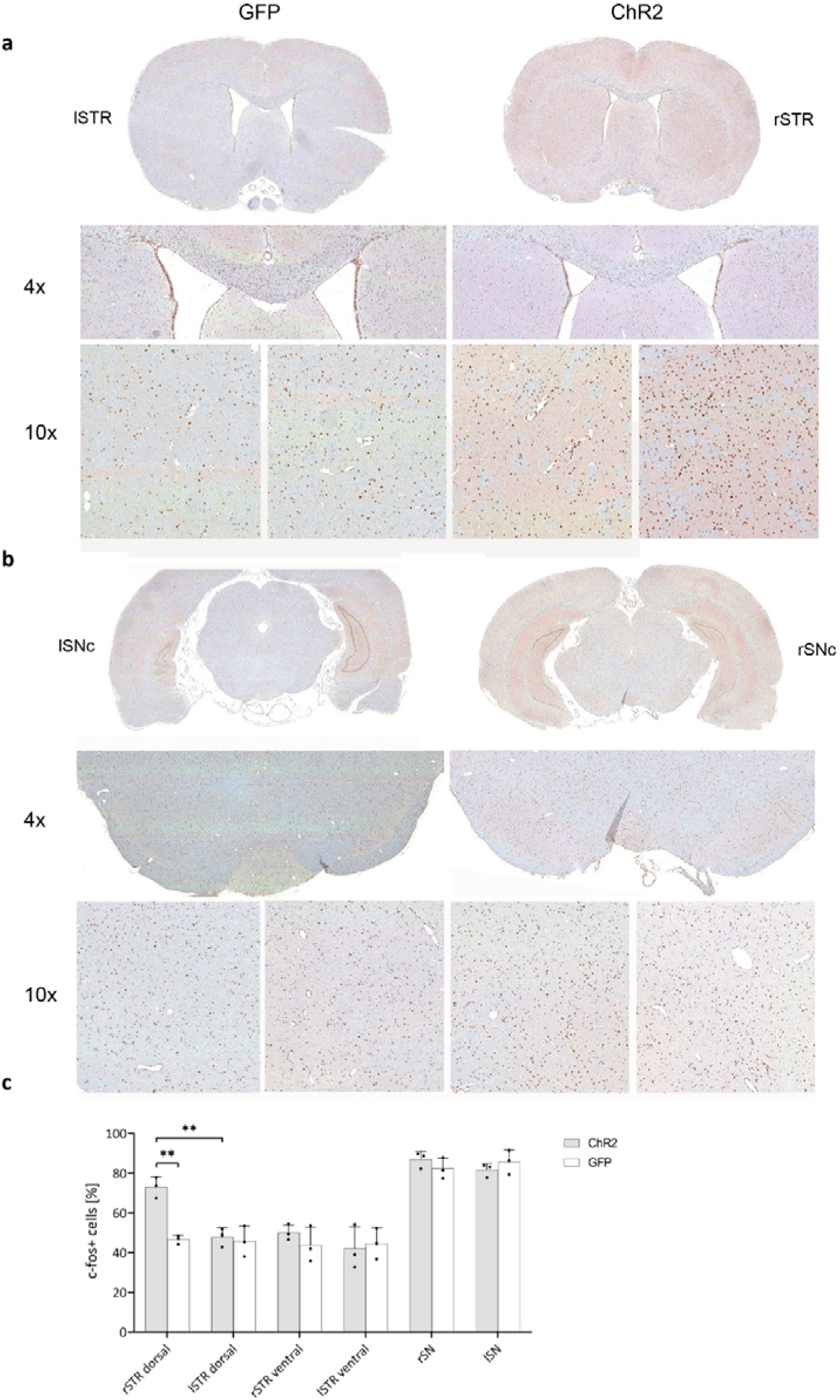
C-fos immunohistochemical staining with quantification in the striatum and substantia nigra. (a) C-fos staining in one exemplary ChR2 and GFP rat is illustrated for one selected region of the dorsal striatum in 1×, 4× and 10× magnifications. A higher number of c-fos+ cells was identified in the 4× and 10× magnifications of the right striatum of the exemplary ChR2 rat. (b) C-fos staining in one exemplary ChR2 and GFP rat is illustrated for one region of interest of the substantia nigra in 1×, 4× and 10× magnifications. (c) The percentage of c-fos+ cells is increased in the right dorsal striatum of the three selected ChR2 expressing rats compared to the left dorsal striatum (*p* = 0.0036) of the ChR2 expressing rats and compared to the right dorsal striatum (*p* = 0.0061) of the three selected GFP expressing rats. Abbreviations: ChR2, channelrhodopsin-2; GFP, green fluorescent protein; l, left; r, right; STR, striatum; SN, substantia nigra

Quantitative analysis revealed a higher percentage of c-fos+ cells in the right dorsal striatum of ChR2 expressing rats compared to the left dorsal striatum (73 ± 4.0% vs. 48 ± 5.6%, *p* = 0.0036) and compared to the right dorsal striatum of GFP expressing rats (47 ± 6.7%, *p* = 0.0061) (see **Supplementary Table 3**, and **Fig. 5c**). No significant differences were observed in the ventral striata and substantia nigra within and between groups.

### Validation of AAV expression

All perfused brains showed normal histology and no pathological alterations were identified. Minor bleedings were focally detected, most likely from the implantation of the optical fiber into the substantia nigra.

ChR2-eYFP and GFP expression in the right striatum and SNc was validated by fluorescence microscopy after staining for GFP/YFP as shown in **Fig. 1b** for one exemplary rat of each group. In some rats, expression of the virus was also observed in the area surrounding the SNc, namely the substantia nigra pars reticulata, the ventral tegmental area and midbrain regions above the substantia nigra pars compacta, which is likely attributed to needle retraction.

### Exclusion of virus-induced neurotoxic effects via TH-staining

The TH IHC in the substantia nigra revealed abundant TH+ neurons and its projections but without qualitative differences between the right and left sides in both GFP and ChR2 expressing rats (**Supplementary Fig. 6**). Additionally, massive presence of TH+ fibers was detected in the striatum, which is known to be a major post-synaptic target of the substantia nigra. In the striatum the TH IHC did not reveal any qualitative differences in the dorsal/ventral and right/left striatum between GFP and ChR2 expressing rats.

## Discussion

fMRI and fPET are two valuable imaging techniques used in neuroscience research to study brain activation. In this study, we employed an optogenetic stimulation of the dopaminergic pathway using simultaneous BOLD-fMRI and [^18^F]FDG-fPET imaging in the rat brain. The findings reveal nuanced insights into both temporal and molecular aspects of brain function.

To enable within-group comparison of fPET data in rats, we employed an [^18^F]FDG bolus+constant infusion protocol. In this study, we demonstrated the application of ICA as a data-driven approach for analyzing rat fPET data acquired during optogenetic stimulation. By automatically sorting components according to spatial kurtosis [37, 38], we identified the [^18^F]FDG uptake map for the optogenetic stimulation component as the first kurtosis-ranked component. This was supported by a GLM-based approach, which identified consistent signal changes, thereby fostering efficient future studies by reducing the need for large control groups.

A distinct discovery of our study was the observation of a small BOLD signal increase during the stimulation and a subsequent overshoot after cessation of the optogenetic stimulation within the striatum, nucleus accumbens and amygdala. These regions are recognized for receiving dopaminergic inputs, leading us to propose a possible role of dopamine for this signal shape. Interestingly, when compared to fPET, only a minimal BOLD response was observed in the SNc in our analysis. To corroborate our observations, we conducted an *ex vivo* analysis of cFos expression in both, the striatum and the SNc. Increased cFos expression levels were evident in the dorsal striatum, which receives input from the SNc, but there were no discernible differences in the SNc itself, confirming our fMRI data. This occurred despite a high metabolic response in the stimulated region, suggesting an active suppression of neuronal firing during optogenetic stimulation. Dopamine release is modulated by numerous neuromodulators [45]. One interpretation posits that optogenetic stimulation triggers the release of dopamine thereby activating dopamine D2 auto- and heteroreceptors and inhibiting further activation-induced dopamine release [46, 47]. This process may curtail the maximum attainable BOLD signal increase during stimulation [17] and may be important to regulate neurotransmitter levels at the synapse. This control is essential for the effective operation of the dopaminergic system. Upon termination of the stimulation, the presynaptic autoinhibition is lifted, leading to the BOLD signal overshoot. Although autoreceptors have been known for many years, the complexity of mammalian central nervous system (CNS) circuits makes it difficult to isolate this mechanism from other neurotransmitter effects. Prior studies have shown that GABA can be co-released from dopaminergic nerve terminals [48], adding to the intricate nature of these interactions.

Alongside the positive BOLD signals, we observed stimulation-induced negative BOLD responses. These were not associated with a relative decrease in [^18^F]FDG in any brain region, a finding consistent with previous research by Stiernman *et al.* [14]. While positive BOLD responses were accompanied by increased glucose metabolism, the authors observed that negative BOLD responses in regions of the default mode network did not show reduced glucose metabolism during a working memory task. Subsequent work demonstrated that this dissociation between negative BOLD response and glucose metabolism is dependent on the corresponding task-positive networks [49]. Negative BOLD responses are considered to be a consequence of increased deoxyhemoglobin concentrations [50, 51]. These increases in deoxyhemoglobin occur during increased oxygen consumption compared to a constant cerebral blood flow or during decreased cerebral blood flow compared to a higher, stable or only slightly reduced oxygen consumption. The precise physiological origin of the negative responses remain debated with theories including the “vascular steel” effect [52, 53], the “vascular sharing” effect [54–56] and regional extremely high oxygen consumption resulting from strong neuronal activation which cannot be balanced by cerebral blood flow increases [53]. Additionally, neurotransmitter release might provoke neurovascular responses which can eventually affect the BOLD signal [57, 58]. A recent study further suggests that opioidergic neurotransmission contributes to negative BOLD-fMRI signals in the striatum [59]. We hypothesize that neurotransmitter and vasoactive effects play a crucial role in the positive and negative responses, but further studies are needed to pinpoint the exact molecular mechanisms.

We further observed metabolic and hemodynamic changes in similar regions, yet differences were found in the spatial extent, location of the regional activation center, proportion of overlapping voxels, and significance of activated regions between the two modalities. For instance in the within-group approach, BOLD-fMRI revealed 15 activated brain regions and 9 regions with a negative BOLD response, while [^18^F]FDG-fPET revealed 16 activated brain regions. Of these regions, 8 showed activation in both modalities, with 9 regions having overlapping voxels. The observed discrepancies between the two modalities can be attributed to their distinct physiological readouts. While [^18^F]FDG is a marker of glucose consumption (metabolic response), the BOLD signal is driven by localized changes in blood flow and blood oxygenation (hemodynamic response). Although several studies indicate highest glucose consumption in neurons at the synaptic level, our data also support high glucose consumption at the soma during active inhibition. In contrast to PET, MRI enables *in vivo* imaging with higher spatial resolution, typically in the range of 0.27×0.27 mm^2^ in plane for the applied EPI-BOLD sequence. However, vascular effects are not confined to the activation site, and larger vessels have a more significant contribution to the BOLD signal. As a result, there is a widespread effect that limits spatial resolution and may result in a mislocalization of activation centers [60].

One notable limitation of our study relates to the utilization of anesthesia, a common element in preclinical imaging studies. Anesthesia has the potential to affect vascular and metabolic responses, which can subsequently lead to alterations in the responsiveness to neuronal stimulations. Although medetomidine anesthesia has been proposed for small animal fMRI experiments in earlier research [61–63] it may significantly elevate blood glucose levels [64, 65], reducing the uptake of [^18^F]FDG into the brain [65, 66]. Isoflurane, often employed in [^18^F]FDG-PET experiments, can also substantially influence the BOLD signal owing to its vasodilatory effect [67–70]. Thus, in an effort to optimize the methodological approach, we selected α-chloralose anesthesia. This choice is based on its suitability for both modalities and its strong functional-metabolic coupling effects, which induce robust fMRI-BOLD activations even after weak stimulations, making it appropriate for [^18^F]FDG-fPET imaging [71–73].

The present study identified pronounced activations in regions within the basal ganglia circuitry, including the striatum, thalamus, and cortex. We also detected increased activation in areas such as the amygdala, septum, hippocampus, periaqueductal gray, and orbitofrontal cortex. These findings might be attributed to the partial stimulation of VTA dopamine neurons [1]. Additionally, our study did not involve the use of TH- or DAT-Cre rats for selective dopaminergic stimulation. This means that optogenetic stimulation may have incidentally extended to areas like the substantia nigra pars reticulata, located beneath the SNc.

In conclusion, our study sheds new light on the intricacies of the dopaminergic pathway, providing novel insights into the relationship between BOLD signals and metabolic responses. By employing simultaneous optogenetic BOLD-fMRI and [^18^F]FDG-fPET imaging, we were able to observe a complex interaction involving both hemodynamic and metabolic processes. The novel findings and the application of cutting-edge techniques in this research offer a roadmap for future investigations into brain function. While the present study unveils the potential of these methods and uncovers new aspects of the neuronal mechanisms, it also emphasizes the need for further detailed research to unravel the complexities of the mammalian CNS circuits. Our results not only enhance the current understanding of the brain’s neurotransmitter systems but also pave the way for more focused and nuanced explorations, especially regarding the interactions between different neurotransmitters and their effect on overall brain functionality.

## Data availability

Codes, raw and processed imaging data will be made available upon request from the principle investigator of the study.

## Code availability

Processing scripts used in the data analysis are available from the corresponding author on request.

## Acknowledgements

We thank the technical assistants at the Werner Siemens Imaging Center (WSIC), University of Tuebingen for their invaluable support during the experiments. We further thank Anna Ohmayer and the Weigelin group at the WSIC for their assistance with the fluorescence microscope. Furthermore, we thank Dr. Julia Mannheim and Dr. Andreas Schmid for their technical support, which ensured the reliable performance of the MRI and PET insert.

Furthermore, we would like to acknowledge the contributions of Dr. Xu-ming Chen and Dr. Xin Yu, who were formerly affiliated at the Max Planck Institute for Biological Cybernetics in Tuebingen, Germany. Their assistance in establishing the optogenetic approach at our institute has been instrumental.

We would like to acknowledge the financial support received for this research project from the following sources: *f*ortüne (internal funding program, University of Tuebingen, to Kristina Herfert), the Carl Zeiss Foundation (to Kristina Herfert), and the Werner Siemens Foundation (to Bernd J. Pichler).

## Author Information

### Authors and Affiliations

**Werner Siemens Imaging Center, Department of Preclinical Imaging and Radiopharmacy, Eberhard Karls University, Roentgenweg 13, 72076 Tuebingen, Germany**

Sabrina Haas, Tudor Ionescu, Fernando Bravo, Gina Dunkel, Laura Kuebler, Bettina Weigelin, Gerald Reischl, Bernd J Pichler, Kristina Herfert

**Cluster of Excellence iFIT (EXC 2180) “Image Guided and Functionally Instructed Tumor Therapies”, Eberhard Karls University, Tuebingen, Germany**

Gerald Reischl, Bernd J Pichler, Leticia Quintanilla-Martinez, Irene Gonzalez-Menendez, Gina Dunkel, Bettina Weigelin

**Institute of Pathology and Neuropathology, Comprehensive Cancer Center, Eberhard Karls University, Tuebingen, Germany**

Irene Gonzalez Menendez, Leticia Quintanilla-Martinez

**Department of Psychiatry and Psychotherapy, Medical University of Vienna, Vienna, Austria**

Andreas Hahn, Rupert Lanzenberger

**Comprehensive Center for Clinical Neurosciences and Mental Health (C3NMH), Medical University of Vienna, Vienna, Austria**

Andreas Hahn, Rupert Lanzenberger

### Contributions

Sabrina Haas – data acquisition and analysis, drafting of the manuscript Fernando Bravo – data analysis, drafting, reviewed manuscript

Tudor Ionescu, Andreas Hahn, Rupert Lanzenberger – data analysis, reviewed manuscript

Gina Dunkel, Irene Gonzalez-Menendez, Leticia Quintanilla-Martinez – data acquisition, analysis

Bettina Weigelin – supervision of microscopy experiments Laura Kuebler – data acquisition, reviewed manuscript Gerald Reischl – tracer synthesis, reviewed manuscript

Andreas Hahn, Rupert Lanzenberger – data analysis, reviewed manuscript

Kristina Herfert – development and conceptual design, supervised experiments, drafting and reviewed manuscript, financial support

Bernd J Pichler – financial support, reviewed manuscript

### Corresponding authors

Correspondence to Kristina Herfert

### Consent for publication

All authors agree with the submitted version of the manuscript. The material submitted for publication has not been previously reported and is not under consideration for publication elsewhere.

## Ethics Declarations

### Ethics approval

All rodent experiments were conducted in compliance with the German animal protection law and with the approval of the local authorities (Regierungspräsidium Tübingen, R6/17).

