## Supplemental Figures and Tables for "Active Suppression of the Nigrostriatal Pathway during Optogenetic Stimulation Revealed by Simultaneous fPET/fMRI"

### Supplementary Figures and Tables

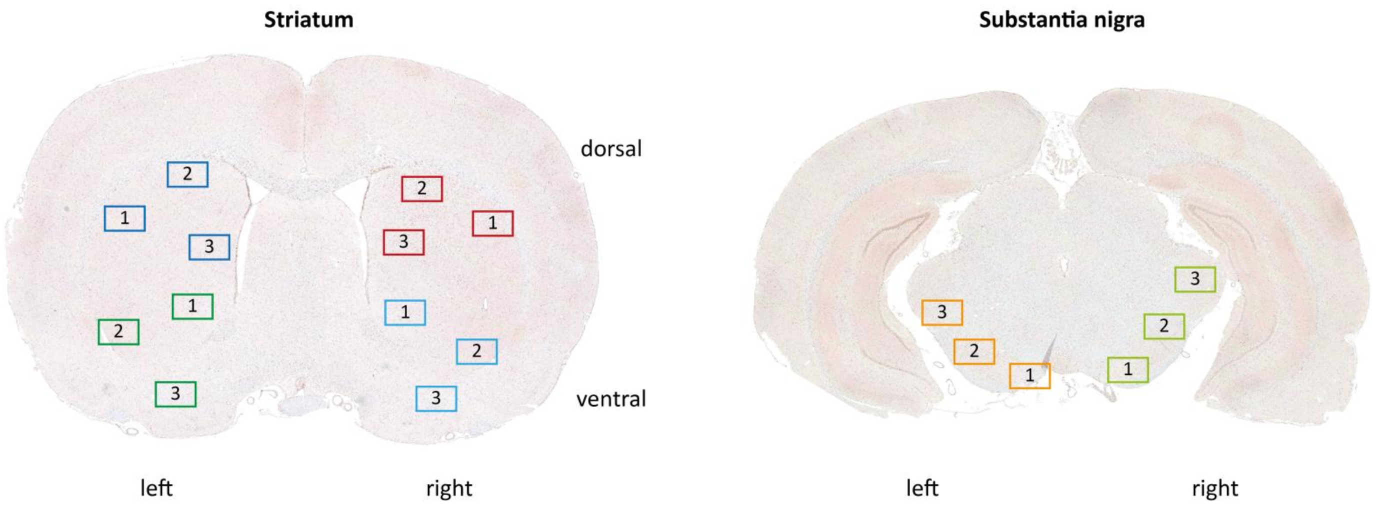


**Supplementary Fig. 1: Selected regions of interest for quantitative analysis of c-fos immunohistochemistry.** Three regions of interest for c-fos quantification were drawn into the right and left, dorsal and ventral striatum and right and left substantia nigra of each selected rat.

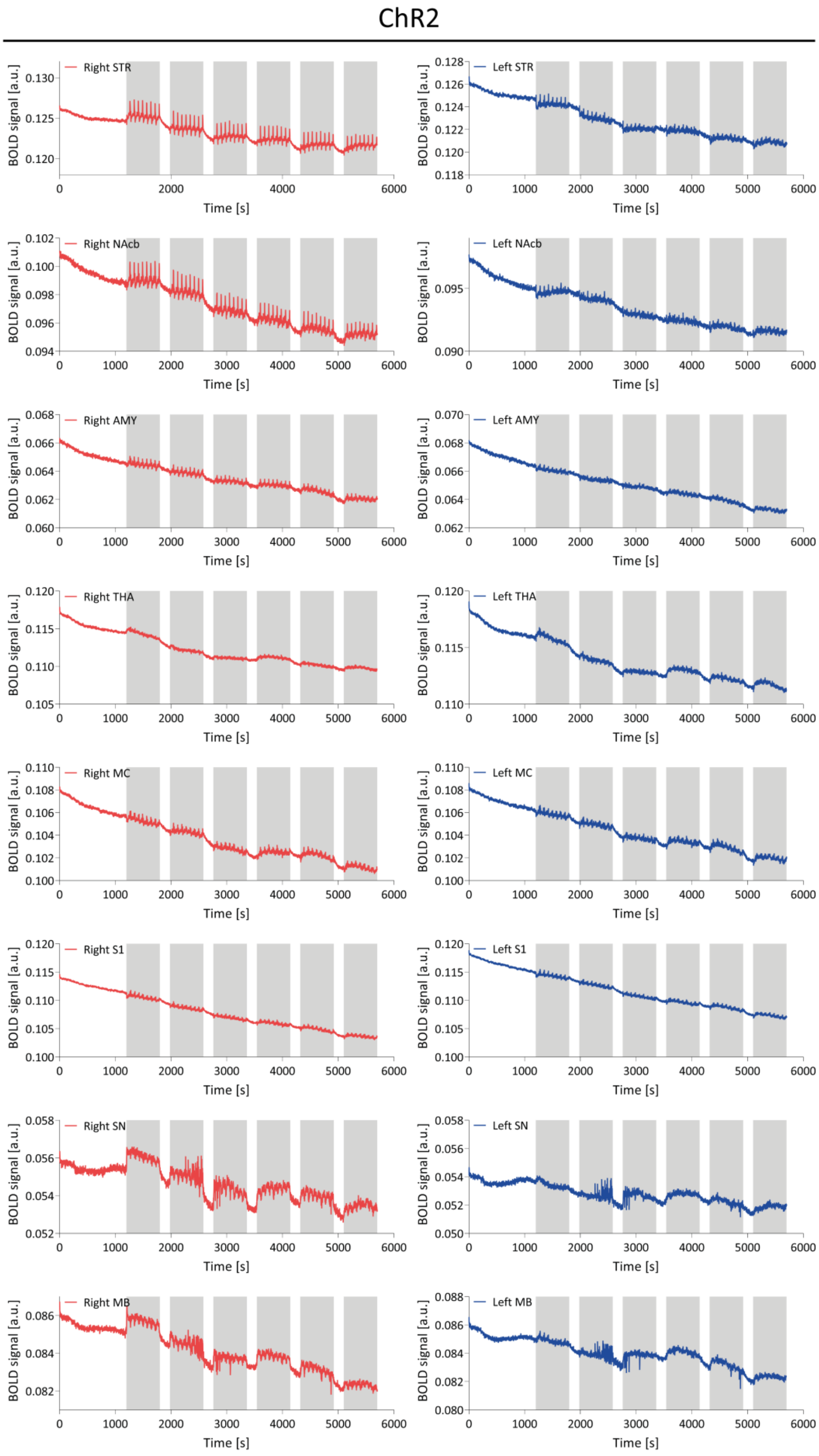


**Supplementary Fig. 2: BOLD-fMRI signal time courses of selected regions.** Mean BOLD signal time courses over 95 minutes are shown for selected regions for ChR2 (n = 18) expressing rats. ChR2 expressing rats responded to stimulation (highlighted in grey) with positive BOLD signal changes in the right striatum, nucleus accumbens, amygdala, thalamus, midbrain and substantia nigra. Negative responses were obtained in the contralateral striatum, nucleus accumbens, amygdala and right and left cortical regions. Abbreviations: AMY, amygdala; MB, midbrain; MC, motor cortex; NAcb, nucleus accumbens; S1, somatosensory cortex; SN, substantia nigra; STR, striatum; THA, thalamus

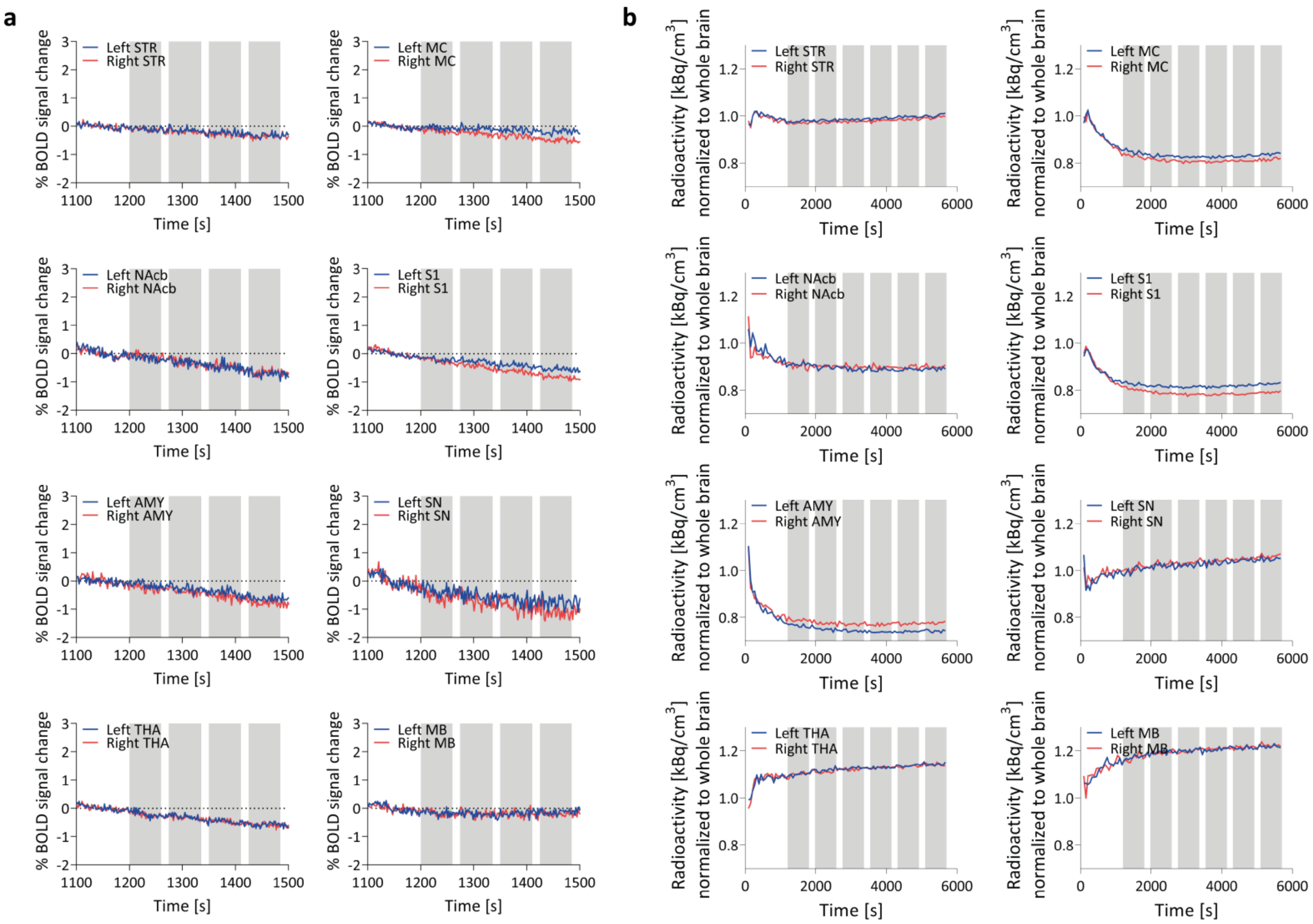


**Supplementary Fig. 3: %BOLD-fMRI signal changes and PET time activity curves in GFP rats.** (**a**) Mean %BOLD signal changes over 400 seconds are shown for selected regions of GFP (n = 12) expressing rats. (**b**) Mean normalized time activity curves over 95 minutes are shown for selected regions of GFP (n = 14) expressing rats. In both modalities, GFP expressing rats did not respond to stimulation (highlighted in grey). Abbreviations: AMY, amygdala; MB, midbrain; MC, motor cortex; NAcb, nucleus accumbens; S1, somatosensory cortex; SN, substantia nigra; STR, striatum; THA, thalamus

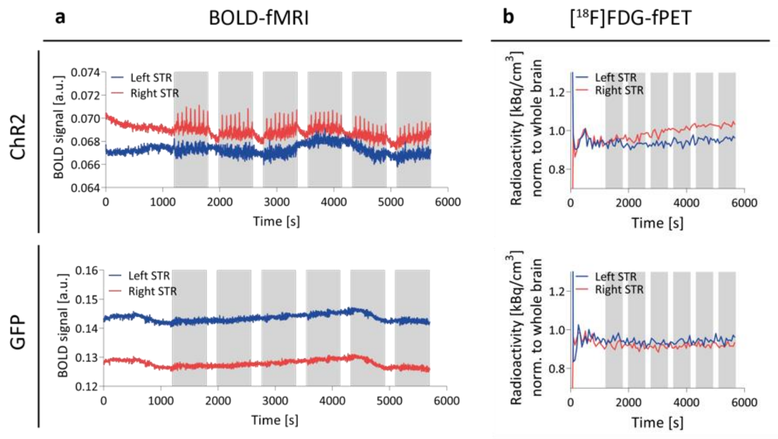


**Supplementary Fig. 4: [$^{18}$F]FDG-fPET and BOLD-fMRI activation on single animal level.** (a) Mean BOLD signal time courses over 95 minutes are shown for the striatum for one exemplary ChR2 and GFP expressing rat. (b) Mean normalized time activity curves over 95 minutes are shown for the striatum of one exemplary ChR2 and GFP expressing rat. Grey bars indicate 10-minute stimulation blocks. Abbreviations: ChR2, channelrhodopsin-2; GFP, green fluorescent protein; STR, striatum

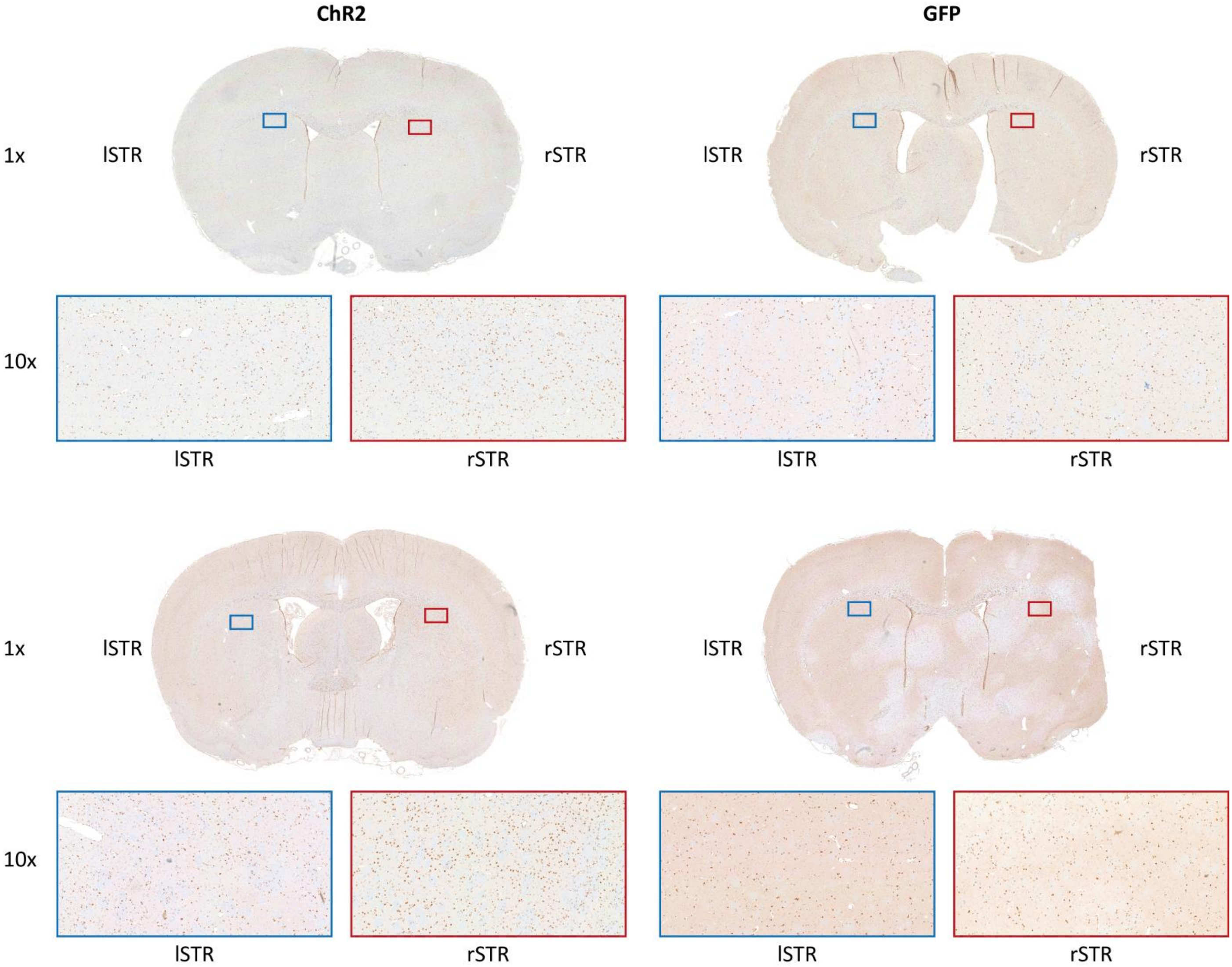


**Supplementary Fig. 5: C-fos immunohistochemical staining in the striatum.** C-fos staining in the other two ChR2 and GFP rats not shown in Fig. 7 is shown for the dorsal striatum in 1× and 10× magnifications. A higher number of c-fos+ cells can be identified in the 10× magnification of the right striatum of the ChR2 rats. Abbreviations: ChR2, channelrhodopsin-2; GFP, green fluorescent protein; l, left; r, right; STR, striatum

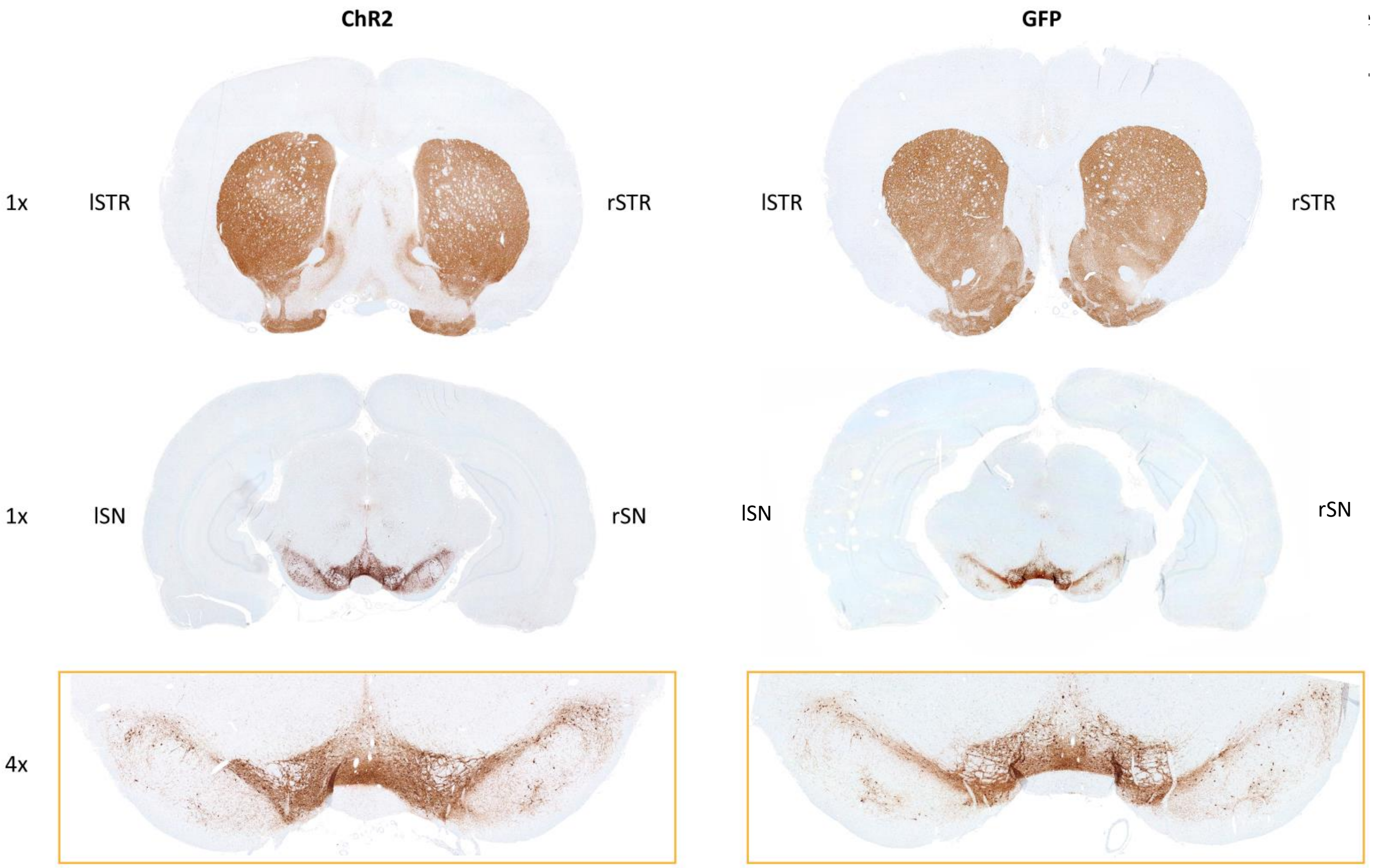


**Supplementary Fig. 6: Tyrosine hydroxylase immunohistochemical staining in the striatum and substantia nigra.** Tyrosine hydroxylase staining in one exemplary ChR2 and one GFP rat is shown for the striatum in 1× and substantia nigra in 1× and 4× magnification. No qualitative left to right differences were identified in neither of the regions. Technical reasons are responsible for an uneven distribution of the staining. Abbreviations: ChR2, channelrhodopsin-2; GFP, green fluorescent protein; l, left; r, right; SN, substantia nigra; STR, striatum

**Supplementary Table 1: Characteristics and abbreviations of selected regions of interest (Schiffer rat brain atlas)**

| Brain region (ROI) | ROI volume [mm³] | # voxels | Abbreviation |
|---|---|---|---|
| **R/ L nucleus accumbens** | 7.9 | 993 | NAcb |
| **R/ L amygdala** | 21.1 | 2640 | AMY |
| **R/ L caudate putamen** | 43.5 | 5444 | STR |
| **R/ L auditory cortex** | 27.5 | 3440 | AC |
| **R/ L cingulate cortex** | 14.5 | 1810 | Cg |
| **R/ L entorhinal cortex** | 59.0 | 7377 | EC |
| **R/ L insular cortex** | 21.1 | 2641 | INS |
| **R/ L medial prefrontal cortex** | 6.3 | 788 | mPFC |
| **R/ L motor cortex** | 32.6 | 4076 | MC |
| **R/ L orbitofrontal cortex** | 18.9 | 2367 | OFC |
| **R/ L parietal cortex** | 7.6 | 954 | PaC |
| **R/ L retrosplenial cortex** | 18.9 | 2365 | RS |
| **R/ L somatosensory cortex** | 71.6 | 8950 | S1 |
| **R/ L visual cortex** | 36.1 | 4517 | V1 |
| **R/ L anterodorsal hippocampus** | 25.1 | 3133 | HIP ant. |
| **R/ L posterior hippocampus** | 9.8 | 1223 | HIP post. |
| **R/ L hypothalamus** | 18.4 | 2294 | HYP |
| **R/ L olfactory cortex** | 14.0 | 1751 | OC |
| **R/ L superior colliculus** | 7.1 | 892 | SC |
| **R/ L midbrain** | 11.4 | 1431 | MB |
| **R/ L ventral tegmental area/ substantia nigra** | 5.5 | 691 | SN |
| **R/ L cerebellum – grey matter** | 75.0 | 9374 | CG |
| **R/ L cerebellum – white matter** | 23.4 | 2938 | CW |
| **R/ L inferior colliculus** | 5.7 | 718 | IC |
| **R/ L thalamus** | 30.7 | 3839 | THA |
| **medulla** | 58.2 | 2944 | Med |
| **periaqueductal gray** | 9.9 | 1238 | PAG |
| **pituitary gland** | 5.8 | 733 | PG |
| **septum** | 9.4 | 1170 | Sep |

Abbreviations: L, left; R, right; ROI, region of interest

**Supplementary Table 2: Independent component analysis (ICA). Descriptive measures derived from the independent component's voxels values distribution for all 20 components (kurtosis sorted)**

| Component | Kurtosis | Skeweness | Variability | Frequency |
|---|---|---|---|---|
| **ICA_9 (OGS)** | 9.4702 | 1.5296 | 0.89649 | 0.0015126 |
| **ICA_7** | 6.1892 | -1.1373 | 0.61559 | 0.0019337 |
| **ICA_5** | 6.0444 | -0.029553 | 0.74643 | 0.0024379 |
| **ICA_1** | 5.9096 | -0.92696 | 0.62911 | 0.0026374 |
| **ICA_3** | 5.5793 | -1.6948 | 0.66834 | 0.0030253 |
| **ICA_6** | 5.1538 | 0.45623 | 0.71338 | 0.0029699 |
| **ICA_4** | 5.0307 | 0.20187 | 0.77642 | 0.002992 |
| **ICA_2** | 4.9093 | 0.3029 | 0.77882 | 0.0029532 |
| **ICA_11** | 4.8252 | 0.46421 | 0.81305 | 0.0027648 |
| **ICA_12** | 4.7691 | 0.42947 | 0.74488 | 0.0031804 |
| **ICA_18** | 4.5608 | -0.071605 | 0.77426 | 0.0026208 |
| **ICA_16** | 4.1498 | 0.35611 | 0.92187 | 0.0031472 |
| **ICA_10** | 3.7893 | 0.17116 | 0.85876 | 0.0032524 |
| **ICA_13** | 3.4033 | 0.043359 | 0.86731 | 0.0030973 |
| **ICA_14** | 2.8282 | 0.045367 | 1.011 | 0.0031305 |
| **ICA_8** | 2.7453 | 0.034452 | 0.87565 | 0.0030862 |
| **ICA_20** | 2.6707 | -0.013394 | 0.77546 | 0.0034574 |
| **ICA_15** | 2.5996 | 0.008139 | 0.80949 | 0.0029699 |
| **ICA_17** | 2.5655 | -0.029454 | 0.77274 | 0.0032912 |
| **ICA_19** | 2.226 | 0.12018 | 0.79925 | 0.0028092 |

**Supplementary Table 3: Percentage of c-fos+ cells**

| | ChR2 | | GFP | |
|---|---|---|---|---|
| **Brain region (ROI)** | **Right** | **Left** | **Right** | **Left** |
| **Dorsal striatum** | 73 ± 4.0% | 48 ± 5.6% | 47 ± 6.7% | 46 ± 2.5% |
| **Ventral striatum** | 50 ± 17% | 42 ± 11% | 44 ± 11% | 45 ± 15% |
| **Substantia nigra** | 87 ± 4.0% | 82 ± 5.5% | 82 ± 5.7% | 86 ± 4.2% |

Abbreviations: ChR2, channelrhodopsin-2; GFP, green fluorescent protein; ROI, region of interest
